# Epitranscriptomic rRNA fingerprinting reveals tissue-of-origin and tumor-specific signatures

**DOI:** 10.1101/2024.10.03.616461

**Authors:** Ivan Milenkovic, Sonia Cruciani, Laia Llovera, Morghan C Lucas, Rebeca Medina, Cornelius Pauli, Daniel Heid, Thomas Muley, Marc A. Schneider, Laura V. Klotz, Michael Allgäuer, Ruben Lattuca, Denis LJ LaFontaine, Carsten Müller-Tidow, Eva Maria Novoa

**Affiliations:** Centre for Genomic Regulation (CRG), The Barcelona Institute of Science and Technology, Dr. Aiguader 88, Barcelona 08003, Spain; Universitat Pompeu Fabra (UPF), Barcelona, Spain; Department of Internal Medicine V, Heidelberg University Hospital, Heidelberg, Germany; Molecular Medicine Partnership Unit (MMPU), European Molecular Biology Laboratory (EMBL), Heidelberg, Germany; Division of Mechanisms Regulation Gene Expression, German Cancer Research Center (DKFZ), Heidelberg, Germany; Translational Lung Research Center (TLRC-H), Member of the German Center for Lung Research (DZL), Heidelberg, Germany; Translational Research Unit and Lung Biobank Heidelberg, Thoraxklinik at Heidelberg University Hospital, Heidelberg, Germany; Department of Surgery, Thoraxklinik at Heidelberg University Hospital, Heidelberg, Germany; Institute of Pathology, Heidelberg University Hospital, Heidelberg, Germany; RNA Molecular Biology, Fonds de la Recherche Scientifique (F.R.S./FNRS), Université libre de Bruxelles (ULB), Biopark campus, B-6041 Gosselies, Belgium

## Abstract

Mammalian ribosomal RNA (rRNA) molecules are highly abundant RNAs, decorated with over 220 rRNA modifications. Previous works have shown that some rRNA modification types can be dynamically regulated; however, how and when the mammalian rRNA modification landscape is remodeled remains largely unexplored. Here, we employ direct RNA nanopore sequencing to chart the human and mouse rRNA epitranscriptome across tissues, developmental stages, cell types and disease. Our analyses reveal multiple rRNA sites that are differentially modified in a tissue- and/or developmental stage-specific manner, including previously unannotated modified sites. We demonstrate that rRNA modification patterns can be used for tissue and cell type identification, which we hereby term ‘epitranscriptomic fingerprinting’. We then explore rRNA modification patterns in normal-tumor matched samples from lung cancer patients, finding that epitranscriptomic fingerprinting accurately classifies clinical samples into normal and tumor groups from only 250 reads per sample, demonstrating the potential of rRNA modifications as diagnostic biomarkers.

## INTRODUCTION

Ribosomes are supramolecular complexes responsible for protein synthesis in all domains of life. Each ribosome consists of a small and a large subunit, and is composed of ribosomal proteins (RP) and ribosomal RNA (rRNA). rRNAs are found at the functional core of the ribosome, playing key roles in mRNA decoding and amino acid polymerization ^1^. Eukaryotic ribosomes typically consist of four different rRNA molecules: 18S rRNA, found in the small ribosomal subunit (40S), and 5S, 5.8S and 28S rRNA, found in the large ribosomal subunit (60S) ^2^.

Ribosomal RNA molecules are extensively modified. In particular, in human and murine ribosomes, the 18S and 28S rRNAs harbor more than 220 modifications ^3^ that can alter physico-chemical properties of the rRNA. Consequently, the presence of rRNA modifications can tweak ribosome function, resulting in enhanced protein synthesis fidelity ^4–6^, among other features. On the other hand, loss of specific rRNA modifications can lead to aberrations in ribosome function ^7^ and decreased translation fidelity ^8^. In the human ribosome, 11 different RNA modification types have been described to date, the most abundant being ribose methylation (Nm) and pseudouridylation (Ψ), which are largely deposited by fibrillarin (FBL) ^9,10^ and dyskerin (DKC) ^11^, respectively.

Ribosomes have been historically considered as uniform macromolecular structures that have identical composition across cell types, tissues and conditions. This view, however, has been challenged in the past few years, leading to a change of paradigm in which ribosomes are now surveyed as dynamic entities that can be heterogeneous in their composition ^12^. The heterogeneity of these ‘specialized ribosomes’ can arise from the use of ribosomal protein paralogs ^13–16^, distinct rRNA variants ^17,18^ or differential rRNA modifications ^19,20^, among others ^21,22^. While the rRNA modification landscape has been previously characterized in both murine and human ribosomes, only a handful of studies have so far characterized the rRNA modification landscape in the context of different tissues ^23,24^ or developmental stages ^25^, and most studies and rRNA databases ^26,27^ do not take into account the tissue and/or cell type of origin ^28^ in their annotations. Notably, the different types of rRNA modifications are likely interconnected, but detailed maps of all rRNA modification patterns are lacking. Moreover, we lack understanding of the complex landscape of individualized ribosomes, which might constitute an additional layer of heterogeneity.

The long-read sequencing platform developed by Oxford Nanopore Technologies (ONT) has the potential to revolutionize our understanding of the epitranscriptome by enabling direct sequencing of native RNA molecules ^29^. Direct RNA sequencing (DRS) permits investigating the transcriptome without the need for reverse transcription or PCR, and can in principle capture any RNA modification that is present in the RNA molecules ^30–32^. Thus, DRS can detect multiple modification types simultaneously along a given RNA molecule, without losing the sequence context and while preserving isoform information ^33^. Preliminary works have already shown that rRNA modifications can be identified using DRS ^20,32,34–38^ and that rRNA modifications can be altered upon environmental insults such as antibiotic exposure ^20,39^.

Here, we employ DRS to study human and mouse rRNA modification dynamics across tissues, cell types, developmental stages and cancer states. We identify rRNA modification ‘signatures’ that are characteristic and distinct across tissues, cell types and developmental stages, including several previously unannotated rRNA sites, which we orthogonally validate and identify as pseudouridylated residues. Moreover, we show that upon cancer, these signatures vary, thus constituting novel promising biomarkers that could be further exploited by future diagnostic approaches. Notably, most of the top differentially modified sites identified through epitranscriptomic fingerprinting do not overlap with previously annotated rRNA modified sites, stressing the importance of employing an agnostic approach to study rRNA modifications and their dynamics. We introduce the concept of ‘epitranscriptomic fingerprinting’, a novel approach that allows classifying samples based solely on their rRNA modification patterns. We propose that this approach could be used in the future to identify tissue-of-origin of cancer samples, as well as to predict cancer.

## RESULTS

### Nanopore DRS identifies differentially modified rRNA sites across tissues and developmental stages

Previous works have shown that a subset of 2’O-methylated rRNA sites are differentially methylated during zebrafish and mouse development ^23,25^ (for full list of annotated human and mouse rRNA modifications, see **Table S1** ^27,28^). However, whether these observations can be validated via orthogonal methods, and whether additional rRNA modification types –beyond Nm modifications– might be differentially modified during development and across tissues, remains unexplored. To address this gap, we applied direct RNA nanopore sequencing (DRS) to study the dynamics of rRNA modifications in native rRNA molecules from four distinct mouse organs (brain, heart, liver and testis) and across three different developmental time points (embryo - E15.5, newborn - P3, and adult - P70) (see *Methods*), allowing for the detection of virtually all rRNA modification types (**Figure 1A**).

**Figure 1.**
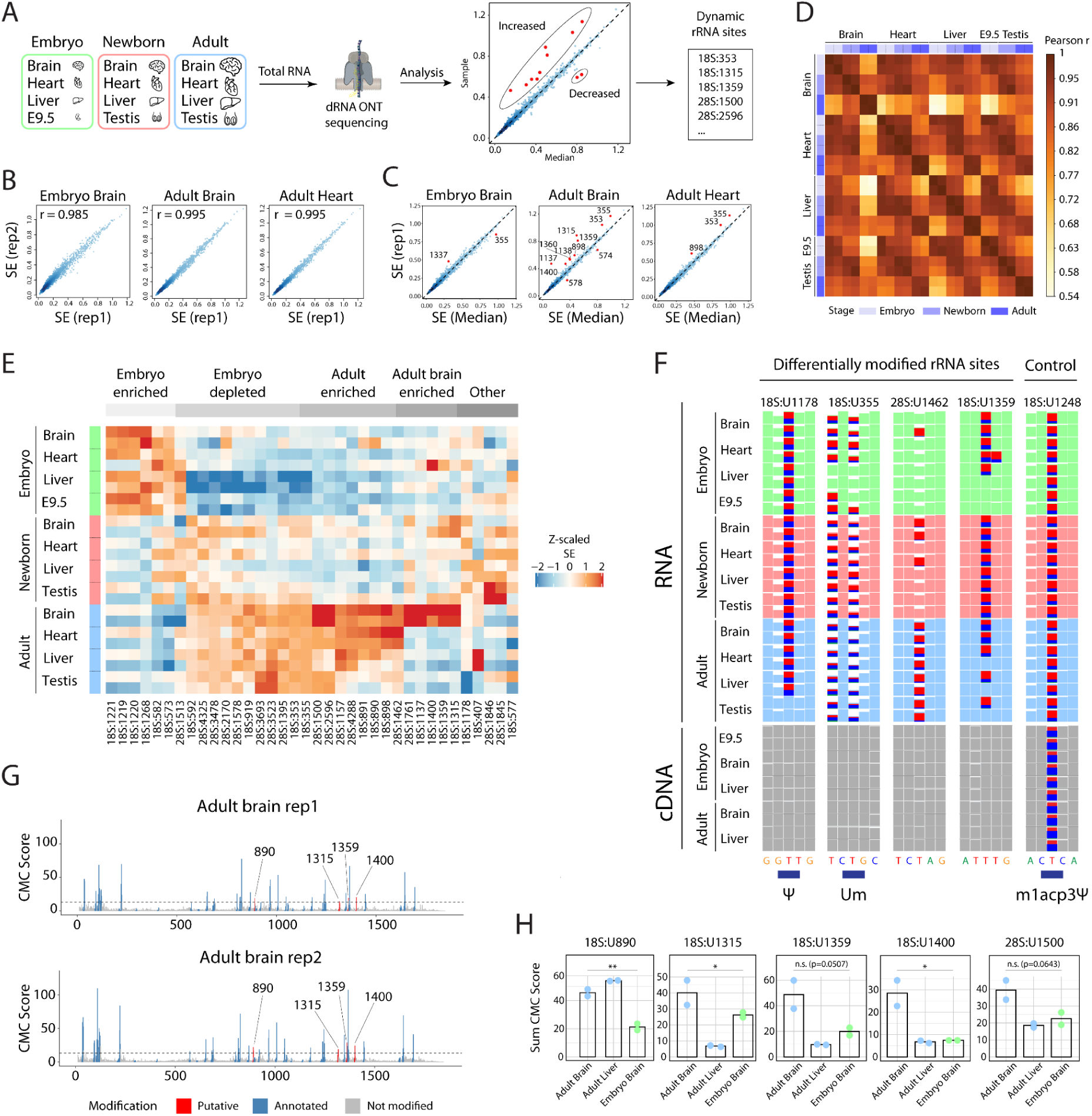
rRNA modification patterns are dynamic across tissues and developmental stages. **(A)** Schematic representation of the workflow used for identification of DM rRNA sites across tissues and developmental stages using direct RNA nanopore sequencing. All experiments were performed on two biological replicates. **(B)** Scatter plots depicting the replicability of SE values computed on 18S rRNA from embryo brain (left), adult brain (middle) and adult heart (right). Pearson r values are shown on the plots. For all replicability plots, see also **Figures S1** and **S2**. **(C)** Scatter plots showing SE value difference from median for embryo brain (left), adult brain (middle) and adult heart (right). Median was calculated from all samples. DM sites (outside the diagonal) are shown in red. For all pairwise comparisons in all samples, see **Figures S3** to **S6**. **(D)** Correlation plots showing Pearson r correlations of SE values calculated on DM sites across all samples for 18S and 28S rRNAs. Developmental stages are shown in the marginal side of the correlation plot, in blue palette colors: embryo (light blue), newborn (blue), adult (dark blue). In the case of testis tissue, embryonic tissue was not collected; E9.5 whole embryo is used at this position in the correlation plot. **(E)** Heatmap of Z-scaled summed error values at DM rRNA sites (including unannotated rRNA modified sites) across tissues and developmental stages. For a heatmap of SE values of all annotated modification sites across tissues and stages, see **Figure S7**. **(F)** IGV snapshot illustrating 4 DM rRNA sites across tissues and/or developmental stages. Two sites correspond to annotated rRNA modified sites (18S:Y1178 and 18S:Um355), whereas the other two (28S:1462 and 18S:1359) are not annotated as rRNA modified sites. The 18S:m^1^acp^3^Ψ1248 site (right) is shown as a example of non–DM site. Embryonic tissues are coloured green, newborn tissues in red and adult tissues in blue. Gray tracks represent cDNA runs from the same samples. Positions with mismatch frequency greater than 0.2 are colored (the proportion of mismatches is shown as stacked histograms with colors corresponding to each base), whereas those showing mismatch frequencies lower than 0.2 are shown in full color (DRS datasets) or gray (cDNA datasets). **(G)** CMC scores along 18s rRNA transcripts, in two independent biological replicates. The CMC score threshold used to define a site as ‘Ψ-modified’ is shown as horizontal dashed line. Known annotated mouse rRNA modified sites are shown in blue, whereas positions that pass the CMC score threshold, but are not annotated as ‘Ψ-modified’, are shown in red (and numerically labeled). See also **Figures S8 and S9** for equivalent tracks in additional tissues and developmental stages. **(H)** CMC scores for the five putative pseudouridylated sites validated by NanoCMC-seq (18S:U890, 18S:U1315, 18S:U1359, 18S:U1400, 28S:1500). CMC scores were calculated by summing the CMC scores of the putatively modified site + 3 downstream positions. One-way ANOVA was used to assess statistical significance of Ψ-modification levels between the sequenced samples (** p.value <0.01; * p.value <0.05; n.s. not significant).

To identify dynamic and/or differentially modified (DM) rRNA sites across tissues and developmental stages, we took advantage of ‘basecalling error’ patterns that are known to occur at modified RNA nucleotides in DRS datasets ^40–43^, to then infer differences (differential basecalling errors) in their modification status across tissues and developmental stages, as previously described ^32^. Specifically, we employed the sum of basecalling errors (mismatches, deletions and insertions), which we refer to as ‘Summed Error’ (SE), and compared the per-site SE values across replicates and conditions. Firstly, we found that SE values were highly robust and replicable between biological replicates (r = 0.977-0.996) (**Figure 1B**, see also **Figures S1** and **S2** and **Tables S2 and S3**). Then, to identify DM rRNA sites, we compared SE values from each sample and site to the median SE value for each site, calculated from all sequenced samples (**Figure 1A**, see also *Methods*). Only those rRNA sites that were identified as differentially modified in both biological replicates were kept for downstream analyses (**Tables S4 and S5**). This approach revealed a total of 31 DM rRNA sites across tissues and/or developmental stages (**Figure 1C**, see also **Figure S3-S6** and **Table S6**). Notably, 14 from the 31 sites identified as differentially modified had not been previously annotated as rRNA-modified sites.

We then systematically compared the SE values for the identified DM rRNA sites (n=31) across replicates, stages and conditions, and computed the Pearson’s correlation for each pairwise comparison (**Figure 1D**). This analysis revealed that the adult brain is the tissue with the most distinct rRNA modification patterns from the set of tissues and stages studied. Afterwards, we examined the SE values at DM rRNA sites, and clustered the samples based on their SE values at these sites (**Figure 1E**). This approach revealed that the set of DM rRNA sites identified could be stratified into: (i) embryo-enriched, (ii) embryo-depleted, (iii) adult-enriched, (iv) adult brain-enriched and (v) other. Visual inspection of rRNA modification patterns at individual sites using IGV confirmed the distinct modification patterns across tissues and stages of these sites (**Figure 1F**).

### Orthogonal validation of previously unannotated rRNA modified sites

To confirm that the DM sites identified here were not caused by SNPs that might be present in some samples but not in others, we also sequenced matched cDNA nanopore sequencing runs from the same samples (**Figure 1F**, shown as bottom gray tracks). Our results show that differential ‘errors’ identified in DRS datasets are not detectable in cDNA tracks from the same samples, indicating that the observed SE variations correspond to rRNA modification differences, and not to SNPs and/or alternative usage of rRNA genes across developmental stages and/or tissues. As a control, we confirmed that we did observe a consistent and reproducible U-to-C mismatch in both the RNA and cDNA reads at the position 18S:1248 which harbors a well-characterized m^1^acp^3^Ψ modification known to affect Watson-Crick base pairing ^44,45^. Overall, our work identified 31 DM rRNA sites, 14 of which had not been even previously annotated as modified rRNA sites, and are hereby regarded as putative rRNA-modified sites (**Table S6**).

To orthogonally validate some of the putative rRNA-modified sites, we performed NanoCMC-seq ^32^ on a subset of samples (adult liver, adult brain and embryo brain), in two independent biological replicates. We were able to validate five of the unannotated putative rRNA modified sites (18S:U890, 18S:U1315, 18S:U1359, 18S:U1400 and 28S:U1500) via NanoCMC-seq, demonstrating that these sites were in fact pseudouridine residues (**Figure 1G**, see also **Figures S8** and **S9**). Moreover, using the semi-quantitative ability of NanoCMC-seq, we confirmed the relative enrichment of these unannotated sites: 18S:Ψ890 was more enriched in adult tissues (both brain and liver), whereas 18S:Ψ1315, 18S:Ψ1359, 18S:Ψ1400 and 28S:Ψ1500 were more enriched in brain, specially in the adult brain (**Figure 1H**), in agreement with our previous observations (**Figure 1E,F**). Finally, we examined whether previously unannotated rRNA modification sites could be assigned to snoRNAs, using snoRNA prediction tools (*see Methods*). These analyses revealed 3 candidate H/ACA box snoRNAs (*Snora35b*, snoRNA-*Taf* and snoRNA-*Wwc1*) possibly guiding 18S:Ψ1315, 18S:Ψ1359 and 28S:Ψ1500 modifications, respectively (**Figure S10,** see also **Table S6).**

### SnoRNA expression levels do not explain differential rRNA modification levels across tissues

We then examined whether the observed differences in rRNA modification levels across tissues and developmental stages identified in this work could be a direct consequence of differences in the snoRNA levels guiding the different modifications. To assess this, we examined publicly available datasets that had quantified small noncoding RNAs across a panel of mouse tissues ^46^.

We then compared the rRNA modification levels with snoRNA expression levels for these 4 snoRNAs across adult brain, heart, liver and testis and found that in this case snoRNA levels do not directly correlate with rRNA modification levels (**Figure S11A**), in agreement with previous studies that had examined this question ^47^. We should note, however, that from the 17 annotated modification sites found to be differentially modified across tissues (**Table S6**), only 4 had a known annotated snoRNA and for which their levels were quantified in the study (namely, Snord90, Snord93, Snord92 and Snora30). Indeed, the top differentially modified rRNA sites that are annotated (18S:Ψ1137, 18S:Ψ1178, 28S:Ψ2596, 18S:Ψ407) do not have as of yet annotated corresponding snoRNAs (**Figure S11B**).

### Dynamic rRNA modification patterns during development can be partially recapitulated *in vitro*

Our analyses point to multiple rRNA modifications as dynamic features that are regulated during development, especially in the brain (**Figure 1D,E**). Considering that cell lines are often used as models to study epitranscriptomic dynamics, we then wondered whether this trend could be recapitulated *in vitro*, by differentiating stem cells into mature neurons. To mimic the three time points used in tissue development (embryo-newborn-adult), we analyzed the rRNA modification patterns of 3 cellular models at different stages of neuronal differentiation: (i) mouse embryonic stem cells (mESC), (ii) neuronal progenitor cells (NPC), and (iii) mature neurons (**Figure 2A**, see also *Methods*). Total RNA was then extracted from each cell type, sequenced using nanopore DRS, and analyzed bioinformatically using the same approach described above (for raw SE values, see **Tables S7 and S8**). We compared DM sites across the brain stages and cell types, finding that in these comparisons mESCs had the most distinct modification profile (**Figure 2B**, see also **Figure S12**). Our analyses showed that rRNA modifications are also differentially modified in *in vitro* neuronal differentiation systems, with some DM rRNA sites during brain development being also mimicked by the *in vitro* system (**Figure 2C**). However, we also found that some modified rRNA sites were only differentially modified *in vitro*, whereas others were only in *in vivo* settings (see **Tables S9** and **S10**). These results suggest that, even though a subset of DM rRNA sites might display similar trends *in vivo* and *in vitro*, findings in these two systems should not be interpreted interchangeably.

**Figure 2.**
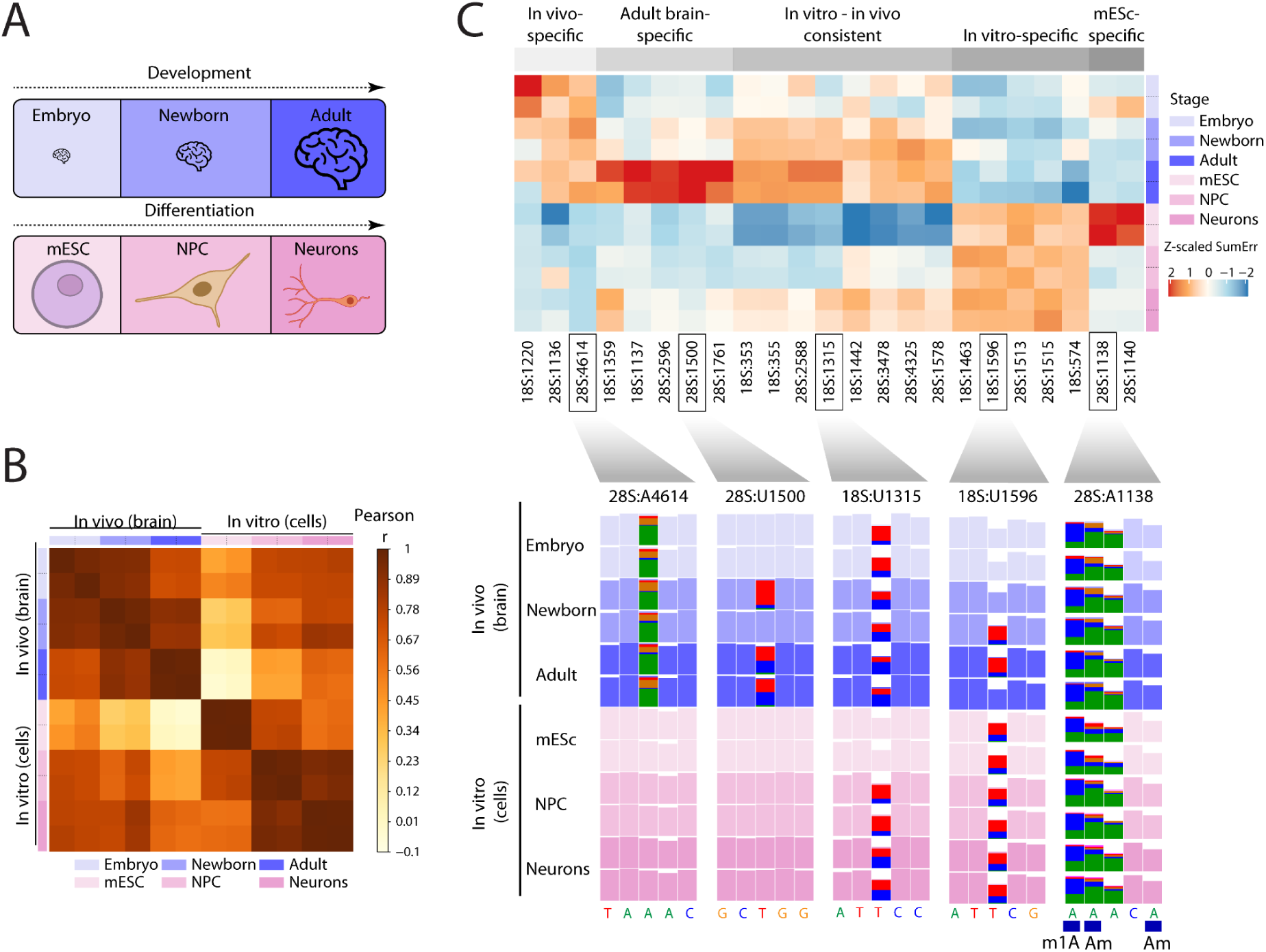
In vitro differentiation models partially recapitulate rRNA modification dynamics observed in the developing brain. **(A)** Experimental design used for in-parallel analysis of rRNA modification profiles from in vivo and in vitro samples. **(B)** Correlation plot showing Pearson r correlations of summed basecalling errors across all samples for 18S and 28S rRNAs, in biological duplicates for each cell type/stage. Developmental stages are shown in the marginal side of the correlation plot, in blue palette colors: embryo (light blue), newborn (blue) and adult (dark blue), while cells are shown in pink palette colors: mESC (light pink), NPC (pink), neurons (dark pink). Biological replicates are shown with the same color. **(C)** Heatmap of summed error values of discovered DM sites (Z-scaled). The discovered DM sites cluster by modification pattern (in vivo-specific, adult brain-specific, in vitro-in vivo consistent, in vitro-specific, mESC-specific). IGV tracks show examples of each type of the discovered sites that are differentially modified. Annotated modification sites are shown below the IGV tracks. Positions with mismatch frequency greater than 0.2 are colored, whereas those showing mismatch frequencies lower than 0.2 are shown in uniform color. For a heatmap of SE values of all annotated modification sites in neuronal development and differentiation, see **Figure S13.**

### rRNA modification patterns accurately predict tissue and developmental stage

Tissue and cell type deconvolution from a given sample are as of yet unsolved problems ^48,49^, especially in the field of cancer research ^50^. A plethora of novel methods have emerged in the last few years to address this issue, most of them typically employing mRNA abundances and/or DNA methylation information as input ^51,52^. Here, we find that rRNA modification patterns are distinct across tissues, cell types and developmental stages (**Figures 1** and **2**). Thus, we wondered whether rRNA modification information captured via DRS could be used for deconvolution of the tissue of origin of a given sample and/or cell type identification.

To examine the potential of rRNA modification information for tissue identification purposes, we first performed an exploratory principal component analysis (PCA) using rRNA modification information from DRS datasets across 4 adult mouse tissues (brain, heart, liver and testis) finding that samples form very distinct clusters based on the tissue type (**Figure 3A**). Notably, pseudouridylated sites were largely responsible for the observed inter-tissue differences (**Figure 3A**, right panel and **Figure 3B**). We then examined whether rRNA modification information would be sufficient to predict tissue type *de novo* from unknown samples based on their rRNA modification patterns (**Figure 3C**). To this end, we randomly subset reads to generate 12 pseudoreplicates for each tissue. The first replicate (flowcell) was used to train the model, and the second replicate (flowcell) was used to test the random forest (RF) model (**Figure S14A**). Afterwards, a third biological replicate was sequenced and used to independently validate the classifier, finding that the trained model could accurately predict all four tissues (accuracy = 1.00, see **Figure S14B**). Similar results were obtained when training classifiers to predict cell types (**Figure S15**) and brain developmental stages (**Figure S16**) from rRNA modification patterns (accuracy = 1.00).

**Figure 3.**
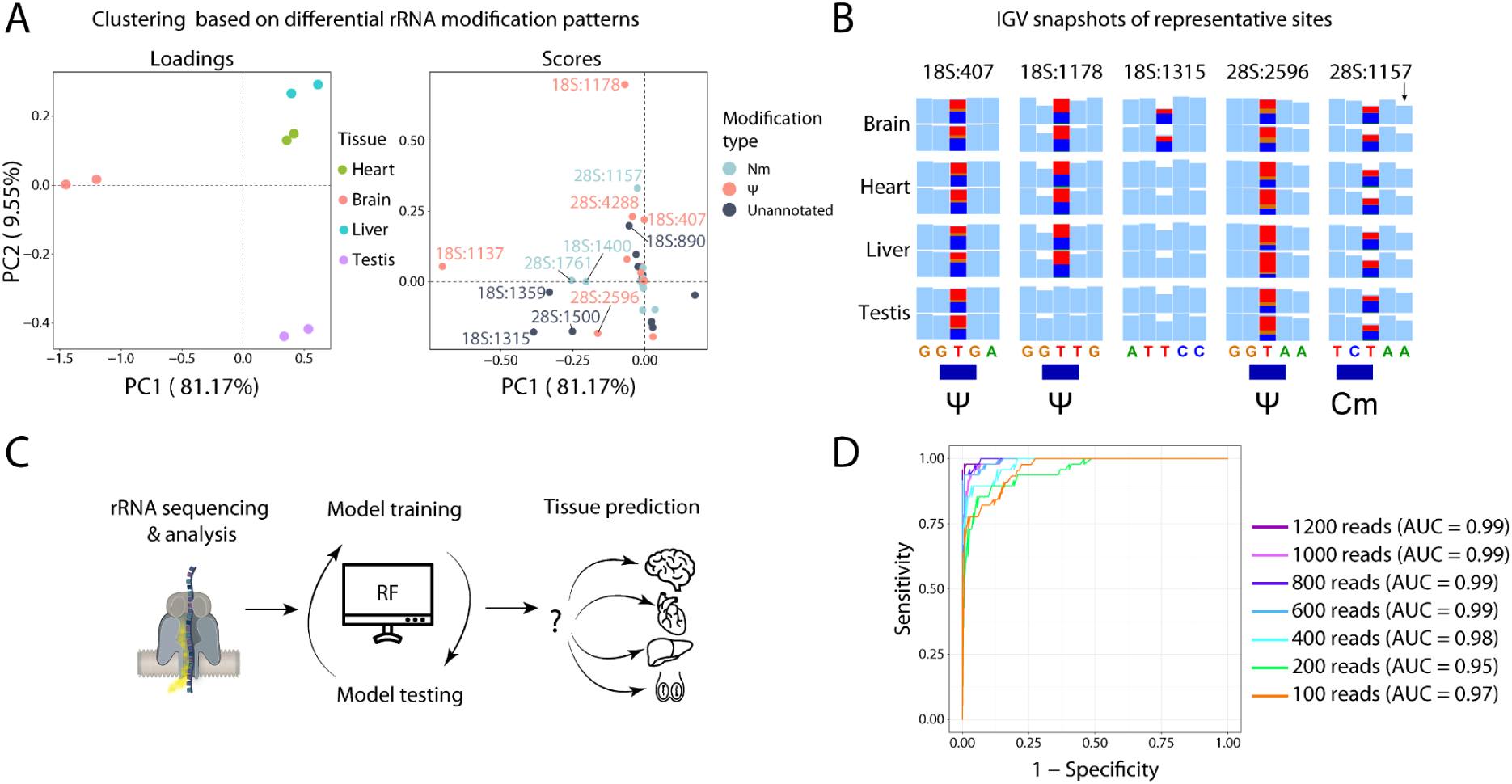
Differential rRNA modification patterns can be used to predict tissue types and developmental stages, using a Random Forest (RF) classifier. **(A)** PCA analysis of adult tissues (brain, heart, liver and testis) for two biological replicates. The scores of DM rRNA sites contributing to the separation of the tissues are shown on the right panel. **(B)** IGV tracks depicting representative examples of tissue-specific DM rRNA sites. Annotated modification sites are shown below the IGV tracks. **(C)** Scheme representing the workflow behind the tissue classifier. First, rRNA is sequenced using DRS and analyzed to obtain information about DM rRNA sites. This information is then used to train a random forest (RF) model, which is then applied to classify tissue types of the testing data set. For more information, see **Figure S14**. **(D)** ROC curves showing the performance of the RF tissue classifier when tested on different numbers of reads (100, 200, 400, 600, 800, 1000 and 1200 reads). For classifiers predicting cell types and developmental stages, see **Figure S15** and **Figure S16**. AUC stands for ‘area under curve’.

Finally, we examined what would be the minimal coverage necessary for accurate tissue classification. To this end, we split the reads from the independent validation DRS dataset into pseudoreplicates with decreasing number of reads (1200, 1000, 800, 600, 400, 200 and 100 reads), finding that even 100 reads were sufficient for the RF classifier to predict tissue types with high accuracy based on their rRNA modification profiles (**Figure 3D**).

### 18S:Um355 ribose methylation is a hallmark of cell proliferation potential, but is not required for neuronal differentiation

We have shown that several rRNA positions displayed consistent and progressive increase in their methylation levels with development (**Figure 1C,F**) as well as upon cellular differentiation (**Figure 2C**). We evaluated whether the modification status of certain rRNA modified sites could be used to assess the proliferation potential of a cell or a tissue sample. To this end, we first performed principal component analysis (PCA) on the discovered DM sites of all tissues and developmental stages (excluding the adult brain, for the full PCA see **Figure S17**), and found that PC1 accurately separated the samples based on their developmental stage (from left to right: adult, newborn and embryo) (**Figure 4A**, left panel). Analysis of the loadings of the PC revealed 18S:Um355 as one of the main contributors for the separation of the samples along the PC1 axis (**Figure 4A**, right panel). Notably, similar results were obtained when performing the same analysis on *in vitro* samples with different proliferative potential (mESC, NPCs and neurons) (**Figure 4B**, left panel), which also showed that 18S:Um355 was one of the main contributing sites for the separation of the cell lines (**Figure 4B**, right panel). Detailed inspection of SE values (proxy of methylation levels) of 18S:Um355 revealed increasing methylation rates matching with cellular differentiation, with mESC having the lowest SE values and adult brain the highest ones (**Figure 4C,D**). Notably, 18S:Um355 has been previously identified as an hypomethylated rRNA modified site in lymphoma ^47^, in agreement with our observations.

**Figure 4.**
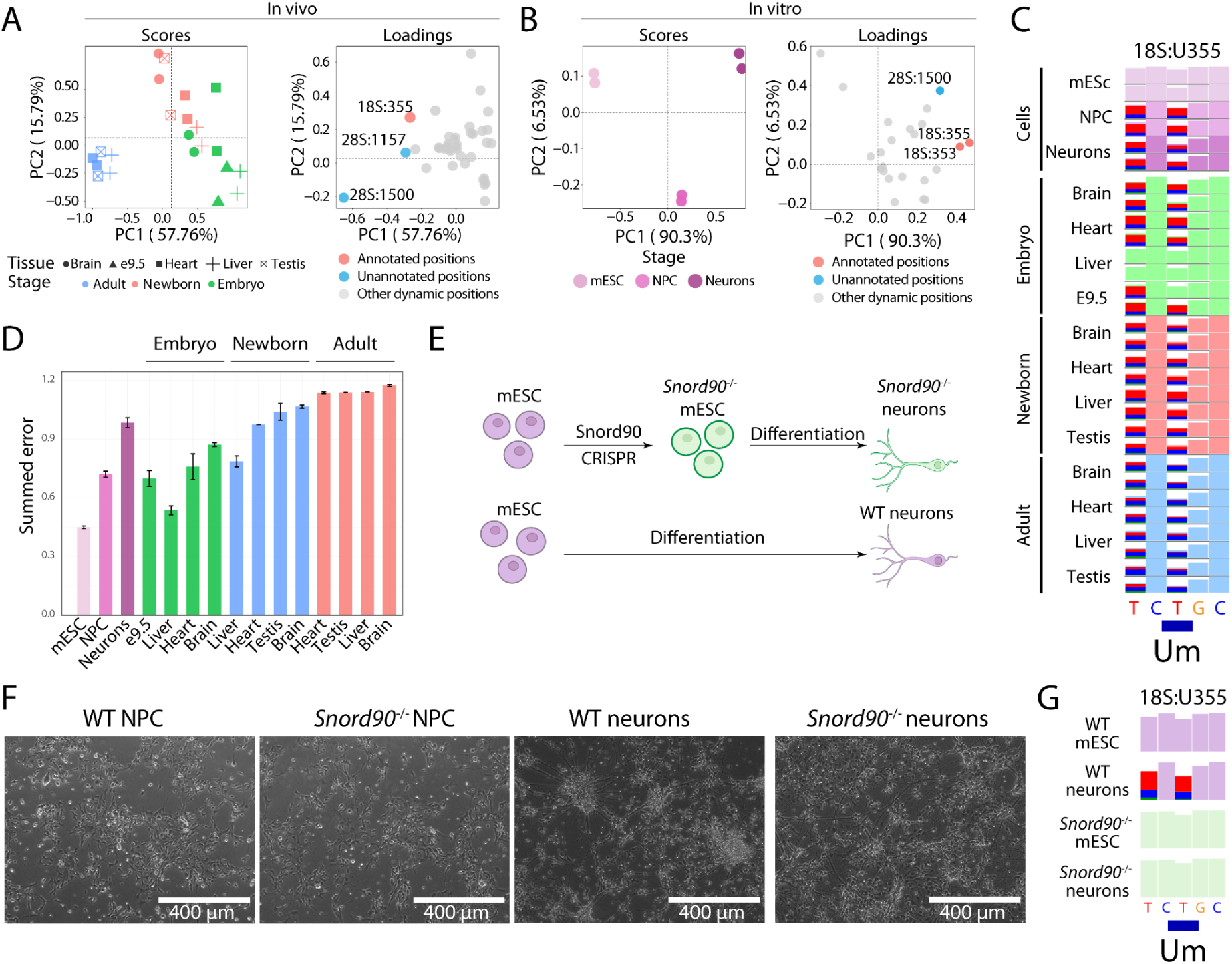
18S:355 methylation status is inversely correlated with the cells’ proliferation potential. **(A)** PCA analysis done on mouse tissue samples (excluding the adult brain). Values used for the PCA are SE of discovered DM sites. **(B)** PCA analysis done on the in vitro samples (mESC, NPCs and Neurons). Values used for the PCA are SE of discovered DM sites. The sites contributing to cluster separation the most are colored red (annotated sites) and blue (unannotated sites); the remaining sites are colored gray (right). **(C)** IGV tracks showing the mismatch pattern of 18S:U355 in mESC, NPC and neuronal cells, as well as in embryonic, newborn and adult tissues. **(D)** Summed error values for 18S:355 for mouse tissue and cell line samples. **(E)** Schematic representation of *Snord90*^−/−^ cell line generation and differentiation to neurons. **(F)** WT and *Snord90*^−/−^ NPCs and neurons. **(G)** IGV tracks showing the mismatch pattern of 18S:U355 in WT and Snord90^−/−^ mESC and neurons. For modification levels of all annotated rRNA sites in WT and *Snord90*^−/−^ mESCs and neurons, see **Figure S18**.

To further dissect the role of 18S:Um355 in differentiation and proliferation, we generated mES cell lines lacking *Snord90*, which is predicted to guide 18S:Um355 modifications ^47^. Because 18S:Um355 is absent in mES (**Figure 4C**), to confirm that Snord90^−/−^ lacked the modification of interest, mES cells were differentiated to NPCs and then to neurons (**Figure 4E,F**, see also *Methods*). The *Snord90^−/−^* NPCs and neurons were indistinguishable from their WT counterparts in terms of cell proliferation, differentiation efficiency and morphology. DRS was performed in the *Snord90*^−/−^ mES and neurons, confirming that the 18S:Um355 mismatch signature was absent in *Snord90*^−/−^-derived neurons (**Figure 4G**, see also **Figure S18**), confirming that snoRD90 is indeed responsible for guiding the methylation at this position. Future work will be needed to assess the role of 18S:Um355 in protein translation.

### Human matched normal and tumor samples exhibit distinct rRNA modification profiles

Previous works have reported that the methylation status (Nm modifications) of some rRNA sites can be dysregulated in cancer ^47,53,54^. Thus, we wondered whether the same could be observed using DRS, and whether additional modifications beyond Nm might be dysregulated in cancer samples. To this end, we sequenced total RNA from matched human normal and tumor tissue samples, namely colon, liver, lung and testis (**Figure 5A**, see also **Table S11** for patient information and **Tables S12** and **S13** for raw summed error values). We discovered that normal and tumor tissues exhibit distinct rRNA modification patterns, especially in the case of lung and testis (**Figure 5B**). PCA analysis on the matched tissues showed that PC1 separated lung and testicular cancer from their matched normal tissues, while colon and liver tumors were harder to distinguish from normal tissues (**Figure 5C**, upper panel). Of note, some of the top DM sites that contributed to the cluster separation (**Figure 5C**, bottom panel) overlapped with sites identified in previous experiments performed on mouse tissues (**Figure 3**). We compared the SE values across tumor and normal tissues (merging different tissues) of the top DM sites, and even though 18S:Ψ1136 was the only significant DM site, we observed a general hypomodification trend in cancer tissues (**Figure 5D**) for all sites. We then performed cDNA sequencing to verify that the basecalling errors were not a product of SNPs but of RNA modifications. Indeed, no basecalling errors were noticeable in the cDNA runs, suggesting that they correspond to rRNA modification differences (**Figure 5E**).

**Figure 5.**
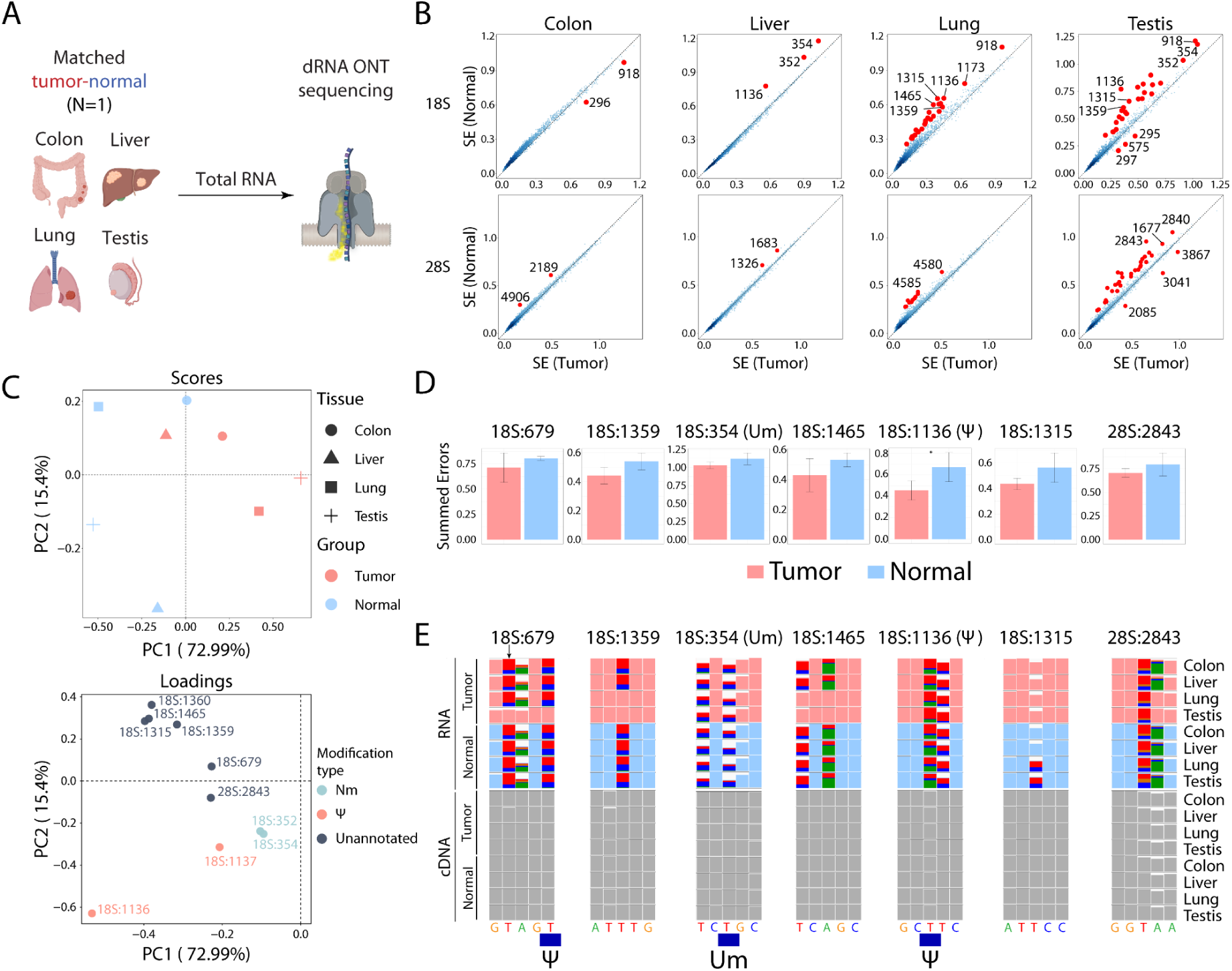
rRNA modification profiles differ between matched normal and human tumor tissues. **(A)** Scheme of tissues used for sequencing. (**B**) SE values for all rRNA sites in 18S (top) and 28S (bottom) in matched colon, liver, lung and testis normal (y-axis) and tumor (x-axis) samples from human donors. The outliers are labeled and shown as red dots. **(C)** PCA analysis performed on matched human samples. The values used for the PCA are discovered DM rRNA sites. The loadings are shown below. **(D)** Average SE values across all cancer and normal samples for the top 7 discovered D; sites. **(E)** IGV tracks of RNA (top) and cDNA (bottom) of matched human cancer and normal tissues for the top 7 discovered DM sites. Annotated modification sites are shown below the IGV tracks, and left blank if no annotated site is present in the shown region.

### rRNA modification profiles can be used to predict the tumor/normal state of matched human lung samples

Following our results from several human tumor and normal tissues, we investigated a larger number of matched tumor-normal human lung tissue samples (N = 20 patients, 40 samples in total; see **Table S14** for patient information and **Tables S15 and S16** for raw SE values). The matched samples were obtained from the same individuals, where normal samples were excised from the same lobe but from a region that was not affected by the tumor (**Figure 6A**, see also *Methods*). Total RNA was extracted and we followed the same workflow as described earlier (**Figure S19A**). Principal component analysis was performed on top DM rRNA sites as previously described, finding a clear separation between normal and tumor samples in the first principal component (PC1) (**Figure 6B**), which explains ~68% of the variance of the data. Notably, our results show that pseudouridylated sites as well as unannotated putative rRNA modified sites are largely responsible for the separation of normal-tumor samples in PC1, with a more modest contribution of Nm-modified sites. Representative IGV snapshots of DM rRNA sites are shown in **Figure 6C**, including some sites that are mostly present in tumor samples (e.g., 18S:Ψ296) and largely hypomodified in normal matched samples, as well as the opposite trend in which the modified site is largely absent or with significantly decreased modification stoichiometry in tumor samples (e.g., 18S:1315), relative to the matched normal samples. We should note that epitranscriptomic fingerprinting could not distinguish between stages I and II (**Figure S20A**), or whether the patients developed metastases or not (**Figure S20B)**.

**Figure 6.**
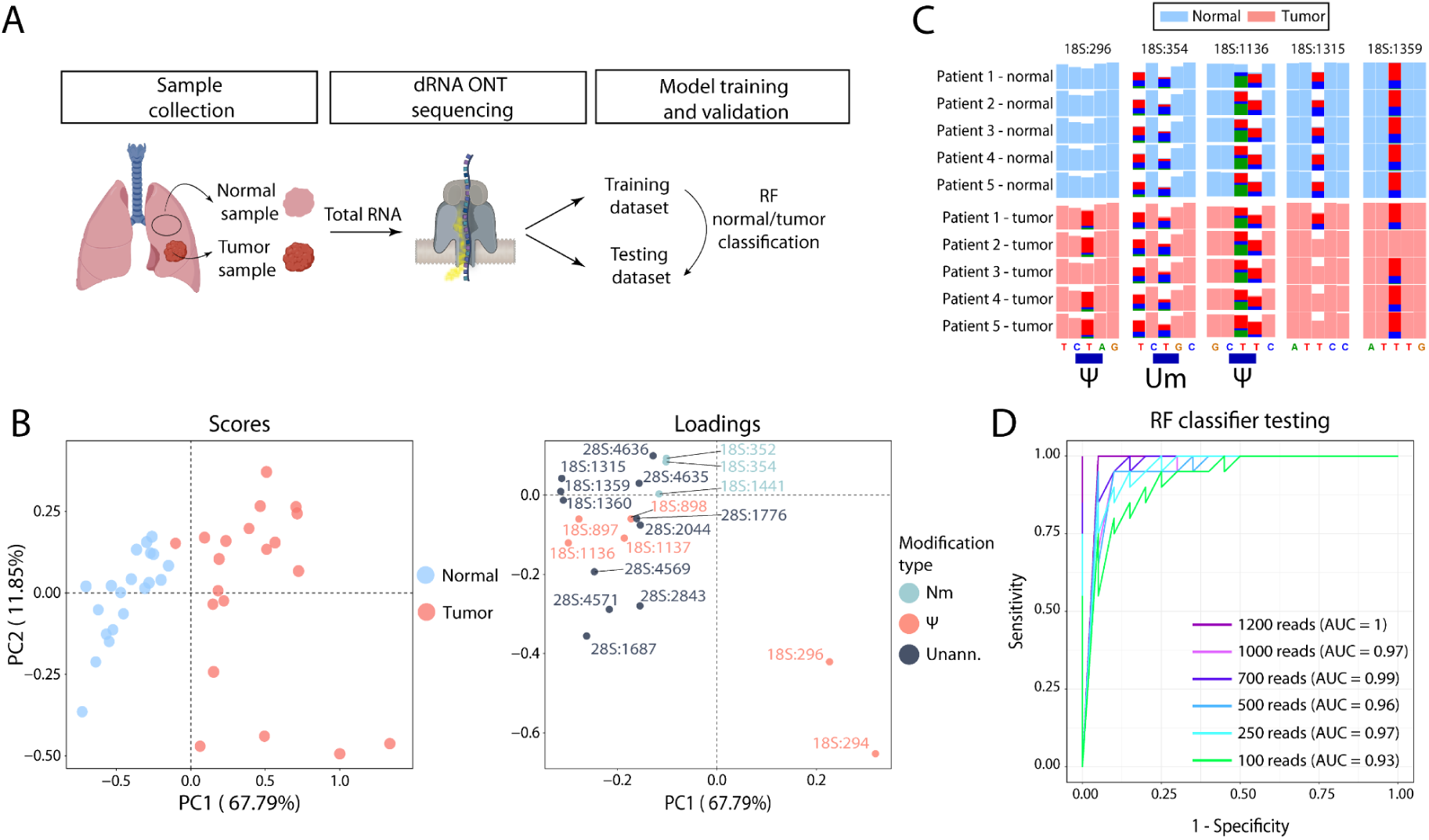
rRNA modification patterns can be used to predict normal/tumor state of lung samples. **(A)** Scheme depicting the sample collection and workflow for training and testing the RF model used for predicting normal/tumor state of lung samples. **(B)** PCA analysis performed on matched lung samples. The values used for the PCA are top 20 discovered DM rRNA sites. The loadings are shown in the bottom panel. **(C)** IGV snapshots of the DM rRNA sites in normal and tumor lung tissues. Annotated modified rRNA sites are shown below the IGV tracks. **(D)** RF normal/tumor classifier performance when tested on different numbers of reads (100, 250, 500, 750, 1000 and 1200 reads). The Area under the curve (AUC) for each coverage threshold is also shown. See also **Figure S19B.**

We then wondered whether the performance trained and tested new RF classifiers using as input only the top 20 DM annotated Nm sites or Ψ sites, and compared the performance of the models with our ‘agnostic’ model that does not require prior knowledge of annotated sites, which employs the top 20 DM sites regardless of the rRNA modification annotations (**Figure S21**). We found that the separation of samples (visualized in their two first principal components) (**Figure S21A**) and classifier performance was significantly decreased when relying only on annotated Nm sites or annotated Ψ-modified sites, compared to using all sites, compared to using all sites –including potentially unannotated ones– (**Figure S21B**).

Finally, we wondered what would be the minimal amount of reads needed for accurate classification of a sample into normal or tumor. To this end, a random forest classifier was trained on the training data set, and internally validated on the test set, and then queried to predict the normal/tumor state of the validation dataset, which had not been used for training/testing the model (**Figure S17,** see also ***Methods***). We randomly subset the testing dataset samples to contain 100, 250, 500, 700, 1000 and 1200 reads, to identify the minimum sequencing depth (number of reads) necessary for successful tumor prediction. After running the classifier on the subsets with different sequencing depths, our results showed that 250 rRNA reads per sample were enough to result in an area of the curve (AUC) of 0.97 (**Figure 6D**). Altogether, our results point to rRNA modifications as promising and powerful biomarkers that could be used to identify disease and/or classify and stratify patient samples in the near future, thus opening a portfolio of novel biomarkers and sequencing methodologies that could be used for cancer identification and/or diagnosis.

## DISCUSSION

In the last few years, the need for cell type identification from sequencing information has emerged ^55^. While mRNAs are typically used for cell type identification and classification ^56^, ribosomal RNAs are routinely discarded from the library preparation and data analysis ^57–59^, as they are generally surveyed as non-informative RNA molecules. In contrast with this view, it has been recently shown that ribosomes, previously thought to be largely invariant, can be heterogeneous in their composition ^13,14,17,60^, including their rRNA modifications ^23,25,61–63^. However, most studies performed to date have largely relied on methods that can only capture one RNA modification type at a time, typically 2’-O-methylations ^23,25,53,62,64^, and have largely limited their analyses to known annotated rRNA sites, thus providing an incomplete picture of rRNA modification dynamics.

Here, we attempted to bridge this gap, and investigated whether rRNA modification patterns can be used as a source of information for tissue-of-origin identification and tumor-normal sample classification. To this end, we employed nanopore DRS to create comprehensive maps of mammalian rRNA modification dynamics across tissues and developmental stages, without restricting the analysis to previously annotated sites or sites with a given RNA modification type. Through this approach, we demonstrated that rRNA modification patterns display tissue-, cell type-, developmental stage- and disease state-specific fingerprints that are easily readable and identifiable using DRS. Notably, we showed that in addition to identifying that some 2’-O-methylations as differentially modified, in agreement with previous works ^23,24^, we found that other rRNA modification types were also differentially modified, with pseudouridylated residues being the most prominent differentially modified rRNA modification type across tissues in all datasets and developmental stages examined (**Figure 3B** and **6B**).

Surprisingly, our analyses also revealed a set of DM rRNA sites at previously unannotated positions (**Figure S22**, see also **Table S17**), suggesting that the list of known rRNA-modified sites is largely incomplete, and that rRNA modifications are not ‘species-specific’, but rather display additional levels of complexity, including tissue-, cell type- and developmental stage-specificity. We then validated five of the putative rRNA modified sites as pseudouridylated sites in mouse using NanoCMC-seq ^32^ (18S:U890, 18S:U1315, 18S:U1359, 18S:1400 and 28S:1500, see **Figure 1G,H** and **Figures S8 and S9**), confirming that the differential basecalling error signatures captured by DRS were indeed coming from pseudouridines that were differentially modified. The differential modification patterns observed using DRS (**Figure 1E**) were partially reproduced by the NanoCMC-seq experiment (**Figure 1H**). Furthermore, we identified three candidate H/ACA box snoRNAs corresponding to three unannotated pseudouridylation sites, one of which is the orphan snoRNA *Snora35b* (**Figure S10, Table S6).** We should point out that 18S:Ψ1315 has also been recently identified in HEK cells ^65^, further supporting our predictions. Altogether, our results not only contribute to completing the picture of rRNA modifications in mouse and human but also highlight the potential of DRS-based approaches for the identification of novel rRNA modifications in previously unexplored biological settings such as in species for which we lack annotated rRNA modifications ^66^.

Our analyses revealed that 18S:Um355 is differentially methylated during development (**Figure 4A**), upon *in vitro* cell differentiation (**Figure 4B**) and in cancer (**Figure 6B**). We propose that the methylation status of 18S:Um355 may be used as a proxy to assess the proliferation rate of cells of origin –with hypomethylation being correlated with high proliferation, and higher methylation levels with more differentiated cells– (**Figure 4A-D**). With regards to the snoRNA guiding 18S:Um355, early works had initially designated this site an ‘orphan’, as it lacked a known specific snoRNA ^27,28^, but more recent works postulated that SNORD90 could be responsible for 18S:355 methylation in human cells ^47^. Subsequent studies overexpressing Snord90 found that this snoRNA could regulate neuregulin 3 (NRG3) mRNA levels, and asserted that Snord90 had no known targets in the rRNA and thus should be considered an orphan snoRNA ^67^. In this work, we demonstrate that SNORD90 is indeed responsible for guiding 18S:Um355 modifications in murine systems, and thus 18S:Um355 should not be further considered as an ‘orphan’ rRNA site (**Figure 4E-G**).

We then examined whether rRNA modification profiles would be altered in cancer, and whether these alterations could be used for sample classification. Notably, previous works had already reported that certain 2’-O-methylated rRNA sites were differentially methylated in specific cancer types ^7,19,21,47,53,68,69^; however, whether other rRNA modification types are differentially modified in cancer remained largely unexplored. We employed DRS to survey the rRNA modification landscape of 20 matched tumor-normal lung cancer human samples (**Figure 6A**), and identified the most variable rRNA sites in terms of modification status. Principal Component Analysis (PCA) showed that most of the variance of the data, explained by PC1, largely separated the samples into two main clusters, which largely correspond to ‘normal’ and ‘tumor’ groups (**Figure 6B**). Our analysis showed that rRNA modification levels typically decrease in cancer samples, relative to normal matched samples, with the exception of 18S:Ψ296, which we found was modified at higher levels in cancer tissues when compared to their corresponding matched normal tissues (**Figure 6B,C**). We should note that previous works have also examined rRNA modification patterns in cancer ^23,25,53,62,64^; however, they largely focused only previously annotated sites, and only one modification type at a time, therefore limiting the amount of new information that could be elucidated. Indeed, we found that top dysregulated sites were in fact unannotated sites, which would have been missed by other approaches restricting the analyses to annotated rRNA modified sites.

Using a Random Forest classifier trained on rRNA modification information, we showed that the trained models can accurately predict tissue (**Figure 3A and S14**), cell type (**Figure S15**) developmental stage (**Figure S16**) and disease state (**Figure 6D** and **S19**), with as few as ~250-500 rRNA reads being sufficient for accurate cell type prediction. Since the average output of a DRS run performed on a MinION flow cell is around 1-2 million reads when using RNA002 kit chemistry (~5M reads with latest RNA004 chemistry)^31,70^, we speculate that dozens to hundreds of samples could be pooled in a single MinION flowcell, once multiplexing of larger number of samples becomes available for DRS^71^, thus greatly decreasing the sequencing costs per sample.

We envision that the ‘epitranscriptomic rRNA fingerprinting’ approach has the potential to be brought to the clinic, for example, applied in the field of early cancer detection. This practice could become especially relevant in the context of screening high-risk populations for diseases such as lung cancer, in which patients are typically diagnosed at late stages (III-IV), often making the disease intractable by surgery or other medical approaches. Indeed, our work demonstrates that rRNA modification information is sufficient to classify samples into ‘normal’ or ‘cancer’ already in lung cancer patients diagnosed with stage I and II tumors (**Table S11**), demonstrating that samples from early cancer stages already present altered rRNA modification patterns that can be captured by the ‘epitranscriptomic fingerprinting’, and providing a strong proof-of-concept of the potential of rRNA modifications to become powerful biomarkers for sample classification and cancer diagnosis. On the other hand, epitranscriptomic fingerprinting could also be useful to guide intraoperative decisions in the future. Of note, intraoperative nanopore sequencing of brain tumors has already been reported in the literature, with sequencing results contributing to the diagnosis and impacting surgical decisions in real time ^72,73^. Future work will be needed to disentangle, examine and validate whether epitranscriptomic fingerprinting can in fact be applied to clinical settings.

Paradoxically, the high abundance of rRNA molecules, typically surveyed for decades as a ‘problem’ in RNAseq libraries, may in fact become a major strength for sample and/or cell type classification especially in low-input scenarios, such as plasma samples. It is yet to be determined whether ‘epitranscriptomic rRNA fingerprinting’ may be applicable to plasma samples, in which reads arising from cancer cells are possibly much less abundant and rRNA molecules are highly fragmented.

### Limitations of the study

While our work demonstrates that rRNAs can act as a source of ribosome heterogeneity across cell types, tissues, developmental stages and disease states, the mechanism whereby this diversity is generated, remains unclear. Differential snoRNA expression has been proposed as a potential source to explain this heterogeneity, however, our re-examination of this possibility (**Figure S11)** as well as by previous works ^47,74^, argue against this hypothesis. Alternatively, this heterogeneity could be mediated by uncharacterised RNA modification-related proteins (RMPs), which could be guided by snoRNAs or not, that would show restricted expression patterns, contributing to the final differential modification levels observed across tissues, cell types, developmental stages and disease states.

On the other hand, the functional consequences of ribosomal rRNA modification diversity also remain unclear. Mechanistically, previous works using cellular knockout models have shown that depletion of specific snoRNAs leads to differences in ribosome footprints ^24^, arguing in favor of the “selective translation” ^75,76^ hypothesis. Under this model, specialized ribosomes generated through differential rRNA modification patterns would be responsible for fine-tuning translation efficiency via preferential translation of a given subset of transcripts. Whether these observations, reported for the 28S:U3904 rRNA modification (guided by SNORD52), are generalizable to other differential modified rRNA sites, is yet to be determined.

Finally, it is still unclear what is the minimum tumor cell content required for accurate detection of ‘cancer’ using ‘epitranscriptomic rRNA fingerprinting’. Future work will be needed to further expand this work onto larger cohorts, as well as in different cancer subtypes.

## STAR★Methods

### Lead contact

Further information and requests for resources and reagents should be directed to and will be fulfilled by the lead contact, Eva Maria Novoa.

### Materials availability

This study did not generate new unique reagents.

### Data and code availability

Base-called FAST5 data used in this work has been deposited into ENA, under accession number PRJEB55716 (study ID ERP140645) for the mouse data, while fastq files for both mouse and human data have been deposited into GEO under GSE264668. A comprehensive list of the datasets generated and used in this work is found in **Table S18**. Raw summed error values for mouse tissues and cells are found in **Tables S2-3** and **S7-8,** and **Tables S12-13** and **S15-16** for matched human normal-tumor samples.

The code used for the analysis of the data presented in this work has been deposited in GitHub https://github.com/novoalab/epitranscriptomic_fingerprinting. Any additional information required to reanalyze the data reported in this study is available from the lead contact upon request.

## Experimental model and study participant details

### Cell lines and culture

Mouse Embryonic Stem Cells (mESC E14tg2A) were cultured in 10% FBS mES complete medium: KnockOut™ DMEM (Thermo, #10829018), MEM Non-Essential Amino Acids Solution, 100X (Thermo, #11140050), GlutaMAX™ 100X (Thermo, #35050061), Sodium Pyruvate 100X (Thermo, #11360070), Pen/Strep 100X (Thermo, #15140122), LIF 107U (Millipore, #ESG1107) 10000X, β-mercaptoethanol 1000X (Sigma, #M-7522, 0,22μm filtered) on 0,1% Gelatine (Millipore #ES-006-B) coated 6-well plates (Thermo, #140675).

Mouse Neuronal Progenitor Cells (mNPCs) were derived from the mESCs with Pro-Neural medium 50% DMEM/F12 GlutaMAX (Gibco, #10565018), 50% Neurobasal medium (Gibco, #21103049), 1% B27 without Vitamin A supplement (Gibco, #12587010), 0.5% N2 supplement (Gibco, #17502048), 1% GlutaMAX (Gibco, #35050-061), 1% Pen/Strep (Thermo, #15140-122) supplemented with 100 ng/ml Noggin (PeproTech, #120-10C) and 20 μM SB431542 (Tebu-Bio, #04-0010-05). mNPCs were cultured in Nunclon Delta surface (Thermo, #150628) 12-well plates coated with 0.001% Poly-L-Ornithine (Sigma, #A-004-M) and 2 μg/ml Laminin (Merck, #L2020) in Pro-neural medium supplemented with Non-Essential Amino Acids Solution (Thermo, #11140050), 10 ng/ml bFGF (Stem Cell Technologies, #78003.1) and 10 ng/ml EGF (Thermofisher, #PMG8041). mNPCs were split every 2-3 days with TrypleExpress (Thermofisher, #12605036) for 3 minutes at RT and seeded at ~250.000 cells per well.

### Mouse models

All adult (P70) C57BL6/J mice were euthanized using CO_2_, and newborn (P3) mice using decapitation. All embryonic tissues were collected at day E15.5, except for whole embryos that were collected at day E9.5. Tissues were quickly excised and snap-frozen in liquid nitrogen, and stored at −80°C until further use.

Animal experimentation was carried out in compliance with EU Directive 86/609/EEC and Recommendation 2007/526/EC regarding the protection of animals used for experimental and other scientific purposes, enacted under Spanish law 1201/2005, and also approved by the institutional ethics committee. Mice used in this study were of C57BL/6J strain background, obtained from Charles Rivers (strain #000664). All mice were raised on a defined control diet (Special Diets Services, RM1 (P), 801151) and were housed in cages at a temperature of 22–24°C, had access to food and water *ad libitum* and were maintained on a 12:12 hour light-dark artificial lighting cycle, with lights off at 19:00.

### Human subjects

Total RNA from human matched tumor and normal tissues were purchased from OriGene (CR560965 - colon tumor; CR560964 - colon normal; CR560583 - lung tumor; CR560571 - lung normal; CR562186 - testis tumor; CR562185 - testis normal; CR559730 - liver tumor; CR561794 - liver normal). The matched samples were collected from the same patients, and each tissue type was collected from a different patient. All the normal tissues were characterized as “within normal limits” during pathology verification. Colon tumor sample was characterized as adenocarcinoma of colon, and graded as G2 (moderately differentiated) following the American Joint Committee on Cancer (AJCC) grading system. Lung tumor sample was characterized as adenocarcinoma of lung, and graded as G3 (poorly differentiated). Testicular tumor sample was characterized as seminoma, and the differentiation status was not reported. Liver tumor sample was characterized as hepatocellular, and graded as G3 (poorly differentiated). See also **Table S12**.

For the matched normal-tumor lung samples, we collected RNA from 20 lung adenocarcinoma patients. The tissue samples were provided from fresh frozen tissue by Lungbiobank Heidelberg, member of the tissue bank of the National Center for tumor diseases (NCT), Germany, in accordance with the regulations of the tissue bank and the approval of the ethics committee of Heidelberg university (S-270/2001, S-568/2023). Tissues were snap-frozen within 30 minutes after resection and stored at −80°C until the time of analysis. For nucleic acid isolation 10 - 15 tumor cryosections (10 - 15 m each) were prepared for each patient. The first and the last sections in each series were stained with hematoxylin and eosin (H&E) and were reviewed by an experienced lung pathologist to determine the proportions of viable tumor cells. Only samples with a viable tumor content of ≥ 50% were used for subsequent analyses. All histopathological diagnoses were made according to the 2015 WHO classification for lung cancer by at least two experienced pathologists. Tumor stage was designated according to the 7th edition of the UICC tumor, node, and metastasis. The cohort is described in **Table S13**.

Human lung cancer samples were obtained from Lung Biobank Heidelberg, a member of the Biobanks of the National Center of Tumor diseases Heidelberg. All tumors (and matching lung tissue) were derived from patients with lung adenocarcinoma undergoing primary tumor resection. Tumor cell content was verified before analysis. Approval was obtained by the Ethics Committee of Heidelberg University Hospital (S-568/2023).

Sequencing and analysis of rRNA modifications using DRS in human samples was approved by the Ethics Committee (CEEA) of the Parc de Salut Mar, under the project proposal 2022/10250/I, titled ‘Development and implementation of nanopore native RNA sequencing technologies for the diagnosis and classification of cancer samples’.

## Method details

### Total RNA extraction from mouse tissues

Tissues were homogenized in TRIzol (Life Technologies, 15596018) using the Polytron PT 1200 E hand homogenizer in pulses of 10 seconds at maximum speed until thoroughly homogenized. Aqueous phase containing RNA was separated by adding chloroform and spinning the samples down for 15 minutes at 12.000 x g at 4 °C. Total RNA was precipitated by adding isopropanol, and then washed with 75% ethanol. Purity and concentration were measured using the NanoDrop spectrophotometer.

### Total RNA extraction from human lung tissues

RNA was isolated using the AllPrep RNA/DNA/miRNA Universal Kit (Qiagen, 80224) using the manufacturer’s instructions. Frozen tumor cryosections were homogenized with the TissueLyser mixer-mill disruptor (2 x 2 min, 25 Hz, Qiagen, Hilden, Germany). The quality of total RNA was assessed with an Agilent 2100 Bioanalyzer and Agilent RNA 6000 Nano Kit (Agilent Technologies, Boeblingen, Germany). The median RIN of the investigated cohort was 8.1 (see **Figure S23**).

### Separation of long and short RNA fractions

Approximately 1.5 µg of each RNA extract was mixed with 3.5X Buffer RLT (Qiagen, #79216) and then 1X 70% ethanol was added to the mix. The samples were transferred to an RNeasy Mini Spin Column (Qiagen, #74104) and centrifuged at 8000 x g for 30 s at RT. The column, containing the long RNA fraction, was kept at RT. The flowthrough, containing the short RNA fraction, was then mixed with 0.65X 100% ethanol, loaded on an RNeasy MinElute Spin Column (Qiagen, #74204), and centrifuged at 8000 x g for 30 s at RT. The flowthrough was discarded and the column was washed first with 700 µl of buffer RWT (Qiagen, #1067933), second with 500 µl of buffer RPE (Qiagen, #1018013) and third with 500 µl 80% ethanol. After the three washes, the empty column was centrifuged for 5 min at 8000 x g with the lid open to remove any residual ethanol. Finally, the short RNA fraction was eluted with 17 µl of RNase-free water. To recover the long RNA fraction, the RNeasy Mini Spin Column was washed twice with 500 µl of buffer RPE. After washing, the column was centrifuged at full speed for 1 min, and then the long RNA fraction was eluted with 30 µl of RNase-free water. Both long and short RNA fractions were quantified by Qubit Fluorometric Quantitation and the RNA electropherogram was obtained using Agilent 4200 TapeStation.

### Differentiation of mouse NPCs to neurons

For neuronal differentiation, mNPCs at ~500.000 cells per well were induced with neuronal differentiation medium, which consisted in Pro-Neural medium supplemented with Non-Essential Amino Acids Solution (Thermo, #11140050), 10 μM DAPT (Selleck, #S2215), 40 ng/ml BDNF (Stem Cell Technologies, #78005.1) for 7 days, replacing 50% of the medium every 2-3 days. A step-by-step protocol for differentiating mouse NPCs to neurons can be found in protocols.io (dx.doi.org/10.17504/protocols.io.6qpvr4oo2gmk/v1).

### mESC *Snord90* knockout generation

The *Snord90* knockout mES cell line was generated using CRISPR-Cas9 editing. ES-E14TG2a mES cells were cultured in regular 10% FBS + LIF conditions on gelatin coated plates. The guides were designed to target a 38 bp-long sequence in the *Rc3h2* host gene, located in the intron coding for *Snord90*. The guide sequences used for CRISPR editing were the following: gRNA upstream: ATTTCATAGGGCAGATTCTG; gRNA downstream: ATTATGAAATCTGAAGACAC. 200.000 mESCs were electroporated using the Neon Electroporation System with a mix of 1.5 pmol of each gRNA and 0.1 μg Cas9 TrueCut protein (Invitrogen, #A36946). A GFP plasmid was used as electroporation control. 48h later, cells were sorted using the regular sorting conditions: 80 μm nozzle, pressure of 120 psi and DAPI as viability dye on a BD Influx cell sorter. 100 clones were retrieved and PCR screened for the 38 bp deletion using the following primer pairs: forward: TTGTACCCTTCCTGTCTCAGAA, reverse: TTAGCAGGGTCACCAGTTCG. 8 clones were validated as successfully edited, carrying the 38 bp deletion.

### Direct RNA nanopore library preparation

All nanopore sequencing runs were performed using the MinION sequencer (flow cell type: FLO-MIN106, sequencing kit: SQK-RNA002). DNase treatment was performed by mixing 2 µg of total RNA per sample with 5 µL of DNase buffer (Life Technologies, #AM2239), 1 µL of RNaseOUT (Invitrogen, #10777019) and 2 µL of Turbo DNase (Life Technologies, #AM2239) in a total volume of 50 µL. The digestion was done by incubating the samples for 10 minutes at 37 °C. The RNA was cleaned using RNA XP beads (Beckman Coulter, #A63987), resuspended in 20 µL of RNase-free water and quantified using NanoDrop spectrophotometer and Qubit™. Poly(A)-tailing was performed by mixing 1 µg of DNase-treated total RNA with the following reagents: 2 µL of poly(A)-tailing buffer (NEB, #B0276SVIAL), 2 µL of 10 mM ATP (NEB, #B0756AVIAL), 1 µL of E. coli Poly(A) polymerase (NEB, #M0276SVIAL), 0.5 µL of SUPERase•In™ (Life Technologies, #AM2694) in a total volume of 20 µL. The reaction mixtures were incubated for 15 minutes at 37 °C. Poly(A)-tailed RNA was then cleaned using Zymo RNA Clean and Concentrator (Zymo, #R1015) and quantified using NanoDrop spectrophotometer and Qubit™. RNA profiles of DNase-treated and poly(A)-tailed RNA were checked using the Tapestation™. Four samples per flow cell were barcoded by ligating custom-made oligonucleotides by mixing the following reagents: 250 ng of poly(A)-tailed total RNA per sample, 1.5 µL NEBNext Quick Ligation Reaction Buffer, 0.75 µL T4 DNA Ligase, 0.5 µL RNaseOUT (Invitrogen, 10777019) and 0.5 µL of pre-annealed custom-made oligonucleotides. The reaction mixtures were mixed by pipetting and incubated for 10 minutes at room temperature. After the ligation, reverse transcription was performed directly by adding 6.5 µL of RNase-free water, 1 µL of 10 mM dNTPs (Thermo Fisher, #18427013), 4 µL of Maxima RT buffer and 1 µL of Maxima reverse transcriptase (Life Technologies, #EP0751). The samples were mixed by pipetting and incubated for 30 min at 60 °C. The RNA-cDNA hybrids were then cleaned using RNA XP beads (Beckman Coulter, #A63987), resuspended in 5 µL of RNase-free water and placed into a clean 1.5 mL Eppendorf DNA LoBind tube (Eppendorf, 30108051). RNA and cDNA were quantified using Qubit™. 50 ng of each sample was pooled together and mixed with 8 µL of NEBNext Quick Ligation Reaction Buffer (NEB, #B6058S), 6 µL of RNA adaptor (RMX, ONT), 3 µL of RNase-free water, and 3 µL of T4 DNA ligase (NEB, #M2200L) in a total volume of 20 µL. The reaction mixture was mixed by pipetting and incubated for 10 minutes at room temperature. The library was cleaned using RNA XP beads (Beckman Coulter, #A63987), with Wash Buffer (WSB, ONT) in the washing steps, for a total of two washes. The beads were then eluted in 21 µL of Elution Buffer (EB, ONT). RNA and cDNA were quantified using Qubit™. The library was mixed with 17.5 µL of RNase-free water and 37.5 µL RRB buffer (ONT) and loaded onto the flow cell.

### Direct cDNA nanopore library preparation

All Nanopore sequencing runs were performed using the MinION sequencer (flow cell type: FLO-MIN106, sequencing kit: SQK-DCS109, barcoding expansion kit: EXP-NBD104). Standard Oxford Nanopore direct cDNA sequencing protocol (version DCB_9091_v109_revC_04Feb2019) was used to sequence mouse tissue total RNA samples. 100 ng of total RNA per sample was used for the first strand synthesis reaction, mixed with 2.5 µL of VNP (ONT cDNA sequencing kit), 1 µL of 10 mM dNTPs and filled to 7.5 µL with RNase-free water. The mixture was incubated at 65 °C for 5 minutes and then snap cooled on ice. The following reagents were mixed in a separate tube: 4 µL of 5x RT buffer, 1 µL RNaseOUT (Invitrogen™), 1 µL of RNase-free water and 2 µL of Strand-Switching Primer (SSP, ONT cDNA sequencing kit). The tubes were gently mixed by flicking and incubated at 65 °C for 2 minutes. 1 µL of Maxima H Minus Reverse Transcriptase (Life Technologies, EP0751) was added to the reaction mixture, which was mixed by flicking and incubated for 90 minutes at 42 °C, followed by heat inactivation at 85 °C for 5 minutes. RNA was degraded by adding 1 µL of RNase Cocktail Enzyme Mix (ThermoFisher, AM2286) followed by incubation for 10 minutes at 37 °C. DNA cleanup was performed using AMPure XP beads and quantity and quality were assessed using Qubit™ and Tapestation™. The second strand was synthesized by mixing the following reagents: 25 µL of 2x LongAmp Taq Master Mix (NEB, 174M0287S), 2 µL PR2 primer (ONT cDNA sequencing kit), 20 µL reverse-transcribed sample and 3 µL RNase-free water. The reaction mixture was incubated using the following protocol: 94 °C, 1 minute; 50 °C, 1 minute; 65 °C, 15 minutes; 4 °C, hold. Another AMPure XP beads cleanup step was performed, proceeding to the end-prep step by mixing the following reagents: 20 µL cDNA sample, 30 µL RNase-free water, 7 µL Ultra II End-prep reaction buffer (NEB, E7647A), 3 µL Ultra II End-prep enzyme mix (NEB, E76468). The mixture was incubated at 20 °C for 5 minutes and 65 °C for 5 minutes. After another AMPure XP beads cleanup step, the samples were barcoded by mixing the following reagents: 22.5 µL End-prepped cDNA, 2.5 µL native barcode (NB01-NB12, ONT barcode extension kit EXP-NBD104), 25 µL Blunt/TA ligase master mix. The reaction mixture was incubated for 10 minutes at room temperature, and the barcoded samples were cleaned up using AMPure XP beads. The cDNA amounts were measured using Qubit™, and the samples were pooled together in equal ratios, not exceeding 120 ng (200 fmol) as the maximum total amount of barcoded cDNA. The adapter ligation was performed by mixing together 65 µL of the pooled barcoded sample, 5 µL Adapter Mix II (AMII, ONT cDNA sequencing kit), 20 µL NEBNext Quick Ligation Reaction Buffer 5X (NEB, B6058S) and 10 µL Quick T4 DNA Ligase (NEB, M2200L). The reaction mixture was incubated for 10 minutes at room temperature, after which the cDNA was cleaned up using AMPure XP beads and eluted in 13 μL of Elution Buffer (EB, ONT cDNA sequencing kit). The final amount was ~50 ng of cDNA, which was mixed with 37.5 µL Sequencing Buffer (SQB) and 2.5 µL Loading Beads (LB, ONT cDNA sequencing kit) and loaded onto a previously primed MinION R9.4.1 flow cell.

### NanoCMC-seq

CMC treatment, library preparation and nanopore sequencing were performed as described by Begik et al. ^32^. 10 µg of total RNA per sample was incubated in NEBNext Magnesium RNA Fragmentation Module at 94 °C for 1.5 min. The fragmented RNA was then incubated with either 0.3 M CMC dissolved in 100 µl of TEU buffer (50 mM Tris pH 8.5, 4 mM EDTA, 7 M urea) or 100 µl of TEU buffer (no CMC) for 20 min at 37 °C. Reaction was stopped with 100 µl of Buffer A (0.3 M NaOAc and 0.1 mM EDTA, pH 5.6), 700 µl of absolute ethanol and 1 µl of Pellet Paint (Novagen, 69049-3). RNA in the stop solution was chilled on dry ice for 5 min and then centrifuged at maximum speed for 20 min at 4 °C. Supernatant was removed, and the pellet was washed with 70% ethanol. After air drying for a few minutes, the pellet was dissolved in 100 µl of Buffer A and mixed with 300 µl of absolute ethanol and 1 µl of Pellet Paint. After chilling on dry ice for 5 min, the solution was then centrifuged at maximum speed for 20 min at 4 °C. Supernatant was removed, and the pellet was washed with 70% ethanol. After washing, the pellet was air dried and resuspended in 40 µl of 50 mM sodium bicarbonate, pH 10.4, and incubated at 37 °C for 3 h. Then, RNA was mixed with 100 µl of Buffer A, 700 µl of ethanol and 1 µl of Pellet Paint for 1 h at −80 °C. The solution was then centrifuged at maximum speed for 25 min at 4 °C, and the pellet was washed with 70% ethanol and dissolved in 30 µL of water after air drying. Unprobed and probed RNAs were treated with T4 Polynucleotide Kinase (NEB, M0201S) for 30 min at 37 °C, before proceeding with ONT Direct cDNA Sequencing.

Before starting the library preparation, 2 µl of 10 µM reverse transcription primer (Original ONT VNP: 5′-/5Phos/ACTTGCCTGTCGCTCTATCTTCTTTTTTTTTTTTTTTTTTTTVN-3′) and 2 µl of 10 µM complementary oligo (CompA: 5′-GAAGATAGAGCGACAGGCAAGTA-3′) were mixed with 1 µl of 0.1 M Tris pH 7.5, 1 µl of 0.5 M NaCl and 4 µl of water. The mix was incubated at 94 °C for 1 min, and the temperature was ramped down to 25 °C (−0.1 °C s−1) to pre-anneal the oligos. The CMC-treated and untreated samples were poly(A) tailed using E. coli Poly(A) polymerase (New England Biolabs, M0276L) for 20 min at 37 °C. Then, 100 ng of poly(A)-tailed RNA was mixed with 1 µl of pre-annealed VNP + CompA, 1 µl of 10 mM dNTP mix, 4 µl of 5× RT Buffer, 1 µl of RNasin Ribonuclease Inhibitor (Promega, N2511), 1 µl of Maxima H Minus RT (Thermo Fisher Scientific. EP0742) and nuclease-free water up to 20 µl. The reverse transcription mix was incubated at 60 °C for 30 min and inactivated by heating at 85 °C for 5 min before moving onto ice. Next, RNAse Cocktail (Thermo Fisher Scientific, AM2286) was added to the mix to digest the RNA, and the mix was incubated at 37 °C for 10 min. Then, the reaction was cleaned up using 1.2× AMPure XP Beads (Agencourt, A63881). To be able to ligate the sequencing adapters to the first strand, 1 µl of 10 µM CompA was again annealed to the 15-µl cDNA in a tube with 2.25 µl of 0.1 M Tris pH 7.5, 2.25 µl of 0.5 M NaCl and 2 µl of nuclease-free water. The mix was incubated at 94 °C for 1 min, and the temperature was ramped down to 25 °C (−0.1 °C s−1) to anneal the complementary to the first-strand cDNA. Next, 22.5 µl of first-strand cDNA was mixed with 2.5 µl of Native Barcode (EXP-NBD104) and 25 µl of Blunt/TA Ligase Mix (NEB, M0367S) and incubated at room temperature for 10 min. The reaction was cleaned up using 1× AMPure XP beads, and the libraries were pooled into one tube that, finally, contained the 200-fmol library. The pooled library was then ligated to the sequencing adapter (AMII) using Quick T4 DNA Ligase (NEB, M2200S) at room temperature for 10 min, followed by 0.65× AMPure XP Bead cleanup using ABB Buffer for washing. The sample was then eluted in elution buffer and mixed with sequencing buffer and loading beads before loading onto a primed R9.4.1 flow cell.

### Nanopore DRS data analysis

Raw fast5 files were basecalled, demultiplexed and mapped using the mop_preprocess module of MoP2 ^77^. Fasta files used as a reference for rRNA mapping were retrieved from GenBank, are correspond to the following annotations: NC_000074.6 (5S rRNA), NR_003280.2 (5.8S rRNA), NR_003278.3 (18S rRNA) and NR_003279.1 (28S rRNA). All fasta sequences are available in: https://github.com/novoalab/epitranscriptomic_fingerprinting/tree/main/fasta_files. The *mop_preprocess* output was used as input for *mop_mod* analysis, and Epinano was used to obtain summed errors calculated at per-site resolution, for each sample. Briefly, the summed error (SE) values were calculated by summing the mismatch, deletion and insertion frequencies at the per-site level for each position in all rRNA molecules, as the frequency of basecalling errors was reported to correlate with the modification status of a given site ^32,40^. The SE values therefore do not quantitatively reflect the modification status (SE = 1 does not mean 100% modified), but they can be used for comparing modification levels across different samples. These values were used for the production of scatter plots, correlation plots and heatmaps, and the scripts used are available at https://github.com/novoalab/epitranscriptomic_fingerprinting. DM rRNA sites were identified by using the following formula: V = |Xi - Med|, where V is the value representing the absolute SE difference between each sample (Xi) and the median for that specific site across all samples (Med). Identified DM sites were considered as “annotated” if i) the signal matched a previously annotated rRNA modification site, or ii) in the case of ribose methylations, if the signal was found at 1-3 bases away from a previously annotated site. Otherwise, the DM sites were regarded as putative unannotated rRNA modification sites. The allele frequency threshold used in all IGV tracks throughout all figures was 0.2.

### NanoCMC-seq data analysis

Raw fast5 files were basecalled, demultiplexed and mapped using the mop_preprocess module of MoP2 ^77^. Fasta files used as a reference for rRNA mapping were retrieved from GenBank, are correspond to the following annotations: NC_000074.6 (5S rRNA), NR_003280.2 (5.8S rRNA), NR_003278.3 (18S rRNA) and NR_003279.1 (28S rRNA). All fasta sequences are available in: https://github.com/novoalab/epitranscriptomic_fingerprinting/tree/main/fasta_files. A custom script prepared by Begik et al. ^32^ was used to extract RT-drop signatures, and the RT-drop scores were plotted using ggplot2. All scripts used to process nanoCMC-seq data with RT-drop information are available in GitHub (https://github.com/novoalab/yeast_RNA_Mod/tree/master/Analysis/NanoCMCSeq). Notably, due to the 5′ end truncation of the nanopore sequencing reads by ~13 nt, RT-drop positions were shifted by 13 nt to accurately determine the exact RT-drop positions. To identify significant RT-drops in a given transcript, we first computed RT-drop scores at each site, which took the difference in the coverage at a given position (0) relative to the previous position (−1). We then computed the difference (Δ RT drop-off score) in RT-drop scores between CMC-probed and unprobed conditions. Lastly, we normalized the Δ RT drop-off score at each position by the median RT drop-off per transcript, leading to final CMC scores, which can be compared across transcripts. Positions with a CMC score greater than 13 for 18S and 12 for 28S were considered significant—that is, to contain a pseudouridine. We should note that the nanoCMC-seq signal-to-noise ratio is dependent on the coverage of the individual transcript.

### Identification of putative mouse snoRNAs

The snoRNA prediction algorithms used were: Snoscan (version 1.0) ^78^ and SnoGPS (version 2.0)^79^, both with default parameters. To localize predicted snoRNA positions on the mouse genome, reads were aligned with STAR (version 2.4.0f1)^80^ on the mouse reference genome mm39 with default parameters and *--outFilterMultimapNmax 1* to discard multi mapped reads. Snoscan and SnoGPS were used on the following datasets: noncoding RNA-Seq of *Mus musculus* lung tissue (PRJNA967497); long RNAs-seq of *Mus musculus* adult male corpus callosum (PRJNA1098504); lariat-intronic RNAs in the *Mus musculus* cytoplasm (PRJNA479418); and small non-coding RNA-seq of *Mus. musculus* heart tissue (PRJNA686442). The following *Mus musculus* reference sequences were used: 18S rRNA (NR_003278.3) and 28S rRNA (NR_003279.1) from NCBI, and genome *mm39* from UCSC.

### Classification of tissues and cell types using random forest models

Bam files of reads mapping to rRNA were randomly subset to contain a portion of random reads for each sample. These pseudoreplicates were then used for RF models training and testing. In the case of mouse tissues, the bam files corresponding to the first two biological replicates were each subset to 12 pseudoreplicates. The bam files were further processed using epinano, and SE values of DM rRNA modification sites were used for building the RF models. The pseudoreplicates corresponding to the first biological replicate were used to train the RF model, which was then used to predict the tissue types of the testing data set, corresponding to the second biological replicate. The training-testing cycle was repeated twice to improve accuracy of the model. The classifier was then validated on the samples corresponding to the third biological replicate (adult brain, heart, liver and testis, with subsetting the bam files to 1200, 1000, 800, 600, 400, 200 and 100 reads). The code to reproduce the results can be found at https://github.com/novoalab/epitranscriptomic_fingerprinting/blob/main/Figure_3/Figure_3_Panel_D.R. The workflow was replicated for cell type and developmental stage classification, with the difference of only using two biological replicates. The RF model was trained on the training data corresponding to the first biological replicate in a single training cycle, and then tested on the testing data corresponding to the second biological replicate.

### Nanopore human data analysis

Raw fast5 files were basecalled, demultiplexed and mapped using the mop_preprocess module of MoP2 ^77^. Fasta files used as a reference for rRNA mapping are the following: NR_023363.1 (5S rRNA); NR_145821.1 (5.8S rRNA), NR_145820.1 (18S rRNA), NR_003287.4 (28S rRNA), and are available at: https://github.com/novoalab/epitranscriptomic_fingerprinting/tree/main/fasta_files. mop_preprocess output was used as input for mop_mod analysis, through which Epinano was run on each sample and summed errors were calculated at per-site resolution. DM rRNA sites were identified by calculating the average SE in cancer and normal samples, and sorting the rRNA sites by the absolute difference between the averages.

## AUTHOR CONTRIBUTIONS

IM performed most of the wet lab experiments and bioinformatic data analyses included in this work. IM built the figures, with the contribution of EMN. SC cultured the mESC and NPCs, and performed neuronal differentiation experiments. LLL performed RNA extractions, library preparation and nanopore sequencing of the human samples used in this work. MCL contributed to the collection of mouse tissue samples used in this work. RM contributed to building some of the DRS libraries used in this work. LK contributed to the collection of human tissue samples. MA evaluated patient samples for tumor content. MAS, TM, DH, CP, CMT provided RNA samples and clinical patient data from the lung cancer cohort. RL and DLJL made snoRNA predictions for unannotated Y modified sites. EMN conceived and supervised the work. IM and EMN wrote the manuscript, with contributions from all authors.

## ACKNOWLEDGEMENTS

We thank all the members of the Novoa lab for their valuable insights and discussion. IM was supported by “la Caixa” InPhINIT PhD fellowship (LCF/BQ/DI18/11660028) and is currently supported by EMBO YIP ‘Bridging Funds’ programme. SC was supported by “la Caixa” InPhINIT PhD fellowship (LCF/BQ/DI19/11730036) and is currently supported by Centro de Excelencia Severo Ochoa funding. MCL was supported by an FPI Severo-Ochoa fellowship by the Spanish Ministry of Economy, Industry, and Competitiveness (MEIC). This work was supported by the European Union’s Horizon 2020 Research and Innovation Program under the Marie Sklodowska-Curie grant agreement [713673], the Spanish Ministry of Economy, Industry and Competitiveness (MEIC) (PID2021-128193NB-100 to EMN), the ERCEA program (ERC-StG-2021 under grant agreement No 101042103 to EMN), the AECC Scientific Foundation (LAB AECC 2021 to EMN) and the German Center for Lung Research (DZL, grant number 82DZL00402). Research in the lab of DLJL was supported by the Belgian Fonds de la Recherche Scientifique (F.R.S./FNRS), EOS [CD-INFLADIS], Région Wallonne (SPW EER) Win4SpinOff [RIBOGENESIS], the COST action TRANSLACORE (CA21154), and the European Joint Programme on Rare Diseases (EJP-RD) RiboEurope and DBAGeneCure. The CMT lab was supported by Deutsche Forschungsgemeinschaft (MU1328/21-1). We also acknowledge the support of the MEIC to the EMBL partnership, Centro de Excelencia Severo Ochoa and CERCA Programme / Generalitat de Catalunya. We thank the CRG Tissue Engineering Facility for their help in generating the *Snord90^−/−^* mESC line and for performing their corresponding neuronal differentiations.

## REFERENCES

1. Nissen, P., Hansen, J., Ban, N., Moore, P.B., and Steitz, T.A. (2000). The structural basis of ribosome activity in peptide bond synthesis. Science 289, 920–930. 10.1126/science.289.5481.920.

2. Khatter, H., Myasnikov, A.G., Natchiar, S.K., and Klaholz, B.P. (2015). Structure of the human 80S ribosome. Nature 520, 640–645. 10.1038/nature14427.

3. Natchiar, S.K., Myasnikov, A.G., Kratzat, H., Hazemann, I., and Klaholz, B.P. (2017). Visualization of chemical modifications in the human 80S ribosome structure. Nature 551, 472–477. 10.1038/nature24482.

4. King, T.H., Liu, B., McCully, R.R., and Fournier, M.J. (2003). Ribosome structure and activity are altered in cells lacking snoRNPs that form pseudouridines in the peptidyl transferase center. Mol. Cell 11, 425–435. 10.1016/s1097-2765(03)00040-6.

5. Baxter-Roshek, J.L., Petrov, A.N., and Dinman, J.D. (2007). Optimization of ribosome structure and function by rRNA base modification. PLoS One 2, e174. 10.1371/journal.pone.0000174.

6. Liang, X.-H., Liu, Q., and Fournier, M.J. (2009). Loss of rRNA modifications in the decoding center of the ribosome impairs translation and strongly delays pre-rRNA processing. RNA 15, 1716–1728. 10.1261/rna.1724409.

7. Zhou, F., Aroua, N., Liu, Y., Rohde, C., Cheng, J., Wirth, A.-K., Fijalkowska, D., Göllner, S., Lotze, M., Yun, H., et al. (2023). A Dynamic rRNA Ribomethylome Drives Stemness in Acute Myeloid Leukemia. Cancer Discov. 13, 332–347. 10.1158/2159-8290.CD-22-0210.

8. Schosserer, M., Minois, N., Angerer, T.B., Amring, M., Dellago, H., Harreither, E., Calle-Perez, A., Pircher, A., Gerstl, M.P., Pfeifenberger, S., et al. (2015). Methylation of ribosomal RNA by NSUN5 is a conserved mechanism modulating organismal lifespan. Nat. Commun. 6, 1–17. 10.1038/ncomms7158.

9. Kiss-László, Z., Henry, Y., Bachellerie, J.P., Caizergues-Ferrer, M., and Kiss, T. (1996). Site-specific ribose methylation of preribosomal RNA: a novel function for small nucleolar RNAs. Cell 85, 1077–1088. 10.1016/s0092-8674(00)81308-2.

10. Tollervey, D., Lehtonen, H., Jansen, R., Kern, H., and Hurt, E.C. (1993). Temperature-sensitive mutations demonstrate roles for yeast fibrillarin in pre-rRNA processing, pre-rRNA methylation, and ribosome assembly. Cell 72, 443–457. 10.1016/0092-8674(93)90120-f.

11. Heiss, N.S., Knight, S.W., Vulliamy, T.J., Klauck, S.M., Wiemann, S., Mason, P.J., Poustka, A., and Dokal, I. (1998). X-linked dyskeratosis congenita is caused by mutations in a highly conserved gene with putative nucleolar functions. Nat. Genet. 19, 32–38. 10.1038/ng0598-32.

12. Xue, S., and Barna, M. (2012). Specialized ribosomes: a new frontier in gene regulation and organismal biology. Nat. Rev. Mol. Cell Biol. 13, 355–369. 10.1038/nrm3359.

13. Li, H., Huo, Y., He, X., Yao, L., Zhang, H., Cui, Y., Xiao, H., Xie, W., Zhang, D., Wang, Y., et al. (2022). A male germ-cell-specific ribosome controls male fertility. Nature 612, 725–731. 10.1038/s41586-022-05508-0.

14. Milenkovic, I., Santos Vieira, H.G., Lucas, M.C., Ruiz-Orera, J., Patone, G., Kesteven, S., Wu, J., Feneley, M., Espadas, G., Sabidó, E., et al. (2023). Dynamic interplay between RPL3- and RPL3L-containing ribosomes modulates mitochondrial activity in the mammalian heart. Nucleic Acids Res. 10.1093/nar/gkad121.

15. Jiang, L., Li, T., Zhang, X., Zhang, B., Yu, C., Li, Y., Fan, S., Jiang, X., Khan, T., Hao, Q., et al. (2017). RPL10L Is Required for Male Meiotic Division by Compensating for RPL10 during Meiotic Sex Chromosome Inactivation in Mice. Curr. Biol. 27, 1498–1505.e6. 10.1016/j.cub.2017.04.017.

16. Guimaraes, J.C., and Zavolan, M. (2016). Patterns of ribosomal protein expression specify normal and malignant human cells. Genome Biol. 17, 236. 10.1186/s13059-016-1104-z.

17. Parks, M.M., Kurylo, C.M., Dass, R.A., Bojmar, L., Lyden, D., Vincent, C.T., and Blanchard, S.C. (2018). Variant ribosomal RNA alleles are conserved and exhibit tissue-specific expression. Sci Adv 4, eaao0665. 10.1126/sciadv.aao0665.

18. Kurylo, C.M., Parks, M.M., Juette, M.F., Zinshteyn, B., Altman, R.B., Thibado, J.K., Vincent, C.T., and Blanchard, S.C. (2018). Endogenous rRNA Sequence Variation Can Regulate Stress Response Gene Expression and Phenotype. Cell Rep. 25, 236–248.e6. 10.1016/j.celrep.2018.08.093.

19. Jansson, M.D., Häfner, S.J., Altinel, K., Tehler, D., Krogh, N., Jakobsen, E., Andersen, J.V., Andersen, K.L., Schoof, E.M., Ménard, P., et al. (2021). Regulation of translation by site-specific ribosomal RNA methylation. Nat. Struct. Mol. Biol. 28, 889–899. 10.1038/s41594-021-00669-4.

20. Delgado-Tejedor, A., Medina, R., Begik, O., Cozzuto, L., Ponomarenko, J., and Novoa, E.M. (2023). Native RNA nanopore sequencing reveals antibiotic-induced loss of rRNA modifications in the A- and P-sites. bioRxiv, 2023.03.21.533606. 10.1101/2023.03.21.533606.

21. Pauli, C., Liu, Y., Rohde, C., Cui, C., Fijalkowska, D., Gerloff, D., Walter, C., Krijgsveld, J., Dugas, M., Edemir, B., et al. (2020). Site-specific methylation of 18S ribosomal RNA by SNORD42A is required for acute myeloid leukemia cell proliferation. Blood 135, 2059–2070. 10.1182/blood.2019004121.

22. Miller, S.C., MacDonald, C.C., Kellogg, M.K., Karamysheva, Z.N., and Karamyshev, A.L. (2023). Specialized Ribosomes in Health and Disease. Int. J. Mol. Sci. 24. 10.3390/ijms24076334.

23. Hebras, J., Krogh, N., Marty, V., Nielsen, H., and Cavaillé, J. (2020). Developmental changes of rRNA ribose methylations in the mouse. RNA Biol. 17, 150–164. 10.1080/15476286.2019.1670598.

24. Häfner, S.J., Jansson, M.D., Altinel, K., Andersen, K.L., Abay-Nørgaard, Z., Ménard, P., Fontenas, M., Sørensen, D.M., Gay, D.M., Arendrup, F.S., et al. (2023). Ribosomal RNA 2’-O-methylation dynamics impact cell fate decisions. Dev. Cell 58, 1593–1609.e9. 10.1016/j.devcel.2023.06.007.

25. Ramachandran, S., Krogh, N., Jørgensen, T.E., Johansen, S.D., Nielsen, H., and Babiak, I. (2020). The shift from early to late types of ribosomes in zebrafish development involves changes at a subset of rRNA 2’-O-Me sites. RNA 26, 1919–1934. 10.1261/rna.076760.120.

26. Quast, C., Pruesse, E., Yilmaz, P., Gerken, J., Schweer, T., Yarza, P., Peplies, J., and Glöckner, F.O. (2013). The SILVA ribosomal RNA gene database project: improved data processing and web-based tools. Nucleic Acids Res. 41, D590–D596. 10.1093/nar/gks1219.

27. Piekna-Przybylska, D., Decatur, W.A., and Fournier, M.J. (2008). The 3D rRNA modification maps database: with interactive tools for ribosome analysis. Nucleic Acids Res. 36, D178–D183. 10.1093/nar/gkm855.

28. Taoka, M., Nobe, Y., Yamaki, Y., Sato, K., Ishikawa, H., Izumikawa, K., Yamauchi, Y., Hirota, K., Nakayama, H., Takahashi, N., et al. (2018). Landscape of the complete RNA chemical modifications in the human 80S ribosome. Nucleic Acids Res. 46, 9289–9298. 10.1093/nar/gky811.

29. Lucas, M.C., and Novoa, E.M. (2023). Long-read sequencing in the era of epigenomics and epitranscriptomics. Nat. Methods 20, 25–29. 10.1038/s41592-022-01724-8.

30. Jain, M., Abu-Shumays, R., Olsen, H.E., and Akeson, M. (2022). Advances in nanopore direct RNA sequencing. Nat. Methods 19, 1160–1164. 10.1038/s41592-022-01633-w.

31. Garalde, D.R., Snell, E.A., Jachimowicz, D., Heron, A.J., Bruce, M., Lloyd, J., Warland, A., Pantic, N., Admassu, T., Ciccone, J., et al. Highly parallel direct RNA sequencing on an array of nanopores. Preprint, https://doi.org/10.1101/068809 10.1101/068809.

32. Begik, O., Lucas, M.C., Pryszcz, L.P., Ramirez, J.M., Medina, R., Milenkovic, I., Cruciani, S., Liu, H., Vieira, H.G.S., Sas-Chen, A., et al. (2021). Quantitative profiling of pseudouridylation dynamics in native RNAs with nanopore sequencing. Nat. Biotechnol. 39, 1278–1291. 10.1038/s41587-021-00915-6.

33. Aw, J.G.A., Lim, S.W., Wang, J.X., Lambert, F.R.P., Tan, W.T., Shen, Y., Zhang, Y., Kaewsapsak, P., Li, C., Ng, S.B., et al. (2021). Determination of isoform-specific RNA structure with nanopore long reads. Nat. Biotechnol. 39, 336–346. 10.1038/s41587-020-0712-z.

34. Fleming, A.M., Bommisetti, P., Xiao, S., Bandarian, V., and Burrows, C.J. (2023). Direct Nanopore Sequencing for the 17 RNA Modification Types in 36 Locations in the E. coli Ribosome Enables Monitoring of Stress-Dependent Changes. ACS Chem. Biol. 18, 2211–2223. 10.1021/acschembio.3c00166.

35. Smith, A.M., Jain, M., Mulroney, L., Garalde, D.R., and Akeson, M. (2019). Reading canonical and modified nucleobases in 16S ribosomal RNA using nanopore native RNA sequencing. PLoS One 14, e0216709. 10.1371/journal.pone.0216709.

36. Grünberger, F., Jüttner, M., Knüppel, R., Ferreira-Cerca, S., and Grohmann, D. (2023). Nanopore-based RNA sequencing deciphers the formation, processing, and modification steps of rRNA intermediates in archaea. RNA 29, 1255–1273. 10.1261/rna.079636.123.

37. Naarmann-de Vries, I.S., Zorbas, C., Lemsara, A., Piechotta, M., Ernst, F.G.M., Wacheul, L., Lafontaine, D.L.J., and Dieterich, C. (2023). Comprehensive identification of diverse ribosomal RNA modifications by targeted nanopore direct RNA sequencing and JACUSA2. RNA Biol. 20, 652–665. 10.1080/15476286.2023.2248752.

38. Stephenson, W., Razaghi, R., Busan, S., Weeks, K.M., Timp, W., and Smibert, P. (2022). Direct detection of RNA modifications and structure using single-molecule nanopore sequencing. Cell Genom 2. 10.1016/j.xgen.2022.100097.

39. Bailey, A.D., Talkish, J., Ding, H., Igel, H., Duran, A., Mantripragada, S., Paten, B., and Ares, M. (2022). Concerted modification of nucleotides at functional centers of the ribosome revealed by single-molecule RNA modification profiling. Elife 11. 10.7554/eLife.76562.

40. Liu, H., Begik, O., and Novoa, E.M. (2021). EpiNano: Detection of m6A RNA Modifications Using Oxford Nanopore Direct RNA Sequencing. Methods Mol. Biol. 2298, 31–52. 10.1007/978-1-0716-1374-0_3.

41. Furlan, M., Delgado-Tejedor, A., Mulroney, L., Pelizzola, M., Novoa, E.M., and Leonardi, T. (2021). Computational methods for RNA modification detection from nanopore direct RNA sequencing data. RNA Biol., 1–10. 10.1080/15476286.2021.1978215.

42. Piechotta, M., Naarmann-de Vries, I.S., Wang, Q., Altmüller, J., and Dieterich, C. (2022). RNA modification mapping with JACUSA2. Genome Biol. 23, 115. 10.1186/s13059-022-02676-0.

43. Jenjaroenpun, P., Wongsurawat, T., Wadley, T.D., Wassenaar, T.M., Liu, J., Dai, Q., Wanchai, V., Akel, N.S., Jamshidi-Parsian, A., Franco, A.T., et al. (2021). Decoding the epitranscriptional landscape from native RNA sequences. Nucleic Acids Res. 49, e7. 10.1093/nar/gkaa620.

44. Motorin, Y., Muller, S., Behm-Ansmant, I., and Branlant, C. (2007). Identification of modified residues in RNAs by reverse transcription-based methods. Methods Enzymol. 425, 21–53. 10.1016/S0076-6879(07)25002-5.

45. Helm, M., and Motorin, Y. (2017). Detecting RNA modifications in the epitranscriptome: predict and validate. Nat. Rev. Genet. 18, 275–291. 10.1038/nrg.2016.169.

46. Isakova, A., Fehlmann, T., Keller, A., and Quake, S.R. (2020). A mouse tissue atlas of small noncoding RNA. Proc. Natl. Acad. Sci. U. S. A. 117, 25634–25645. 10.1073/pnas.2002277117.

47. Krogh, N., Asmar, F., Côme, C., Munch-Petersen, H.F., Grønbæk, K., and Nielsen, H. (2020). Profiling of ribose methylations in ribosomal RNA from diffuse large B-cell lymphoma patients for evaluation of ribosomes as drug targets. NAR Cancer 2, zcaa035. 10.1093/narcan/zcaa035.

48. Ma, Y., and Zhou, X. (2022). Spatially informed cell-type deconvolution for spatial transcriptomics. Nat. Biotechnol. 40, 1349–1359. 10.1038/s41587-022-01273-7.

49. Li, B., Zhang, W., Guo, C., Xu, H., Li, L., Fang, M., Hu, Y., Zhang, X., Yao, X., Tang, M., et al. (2022). Benchmarking spatial and single-cell transcriptomics integration methods for transcript distribution prediction and cell type deconvolution. Nat. Methods 19, 662–670. 10.1038/s41592-022-01480-9.

50. Li, S., Zeng, W., Ni, X., Liu, Q., Li, W., Stackpole, M.L., Zhou, Y., Gower, A., Krysan, K., Ahuja, P., et al. (2023). Comprehensive tissue deconvolution of cell-free DNA by deep learning for disease diagnosis and monitoring. Proc. Natl. Acad. Sci. U. S. A. 120, e2305236120. 10.1073/pnas.2305236120.

51. Li, R., Liao, B., Wang, B., Dai, C., Liang, X., Tian, G., and Wu, F. (2021). Identification of Tumor Tissue of Origin with RNA-Seq Data and Using Gradient Boosting Strategy. Biomed Res. Int. 2021, 6653793. 10.1155/2021/6653793.

52. Chu, T., Wang, Z., Pe’er, D., and Danko, C.G. (2022). Cell type and gene expression deconvolution with BayesPrism enables Bayesian integrative analysis across bulk and single-cell RNA sequencing in oncology. Nat Cancer 3, 505–517. 10.1038/s43018-022-00356-3.

53. Marcel, V., Kielbassa, J., Marchand, V., Natchiar, K.S., Paraqindes, H., Nguyen Van Long, F., Ayadi, L., Bourguignon-Igel, V., Lo Monaco, P., Monchiet, D., et al. (2020). Ribosomal RNA 2’O-methylation as a novel layer of inter-tumour heterogeneity in breast cancer. NAR Cancer 2, zcaa036. 10.1093/narcan/zcaa036.

54. Peng, H., Chen, B., Wei, W., Guo, S., Han, H., Yang, C., Ma, J., Wang, L., Peng, S., Kuang, M., et al. (2022). N6-methyladenosine (m6A) in 18S rRNA promotes fatty acid metabolism and oncogenic transformation. Nat Metab 4, 1041–1054. 10.1038/s42255-022-00622-9.

55. Shekhar, K., and Menon, V. (2019). Identification of Cell Types from Single-Cell Transcriptomic Data. Methods Mol. Biol. 1935, 45–77. 10.1007/978-1-4939-9057-3_4.

56. Katsetos, C.D., Herman, M.M., and Mörk, S.J. (2003). Class III beta-tubulin in human development and cancer. Cell Motil. Cytoskeleton 55, 77–96. 10.1002/cm.10116.

57. O’Neil, D., Glowatz, H., and Schlumpberger, M. (2013). Ribosomal RNA depletion for efficient use of RNA-seq capacity. Curr. Protoc. Mol. Biol. Chapter 4, Unit 4.19. 10.1002/0471142727.mb0419s103.

58. Bush, S.J., McCulloch, M.E.B., Summers, K.M., Hume, D.A., and Clark, E.L. (2017). Integration of quantitated expression estimates from polyA-selected and rRNA-depleted RNA-seq libraries. BMC Bioinformatics 18, 301. 10.1186/s12859-017-1714-9.

59. Loi, D.S.C., Yu, L., and Wu, A.R. (2021). Effective ribosomal RNA depletion for single-cell total RNA-seq by scDASH. PeerJ 9, e10717. 10.7717/peerj.10717.

60. Locati, M.D., Pagano, J.F.B., Girard, G., Ensink, W.A., van Olst, M., van Leeuwen, S., Nehrdich, U., Spaink, H.P., Rauwerda, H., Jonker, M.J., et al. (2017). Expression of distinct maternal and somatic 5.8S, 18S, and 28S rRNA types during zebrafish development. RNA 23, 1188–1199. 10.1261/rna.061515.117.

61. Babosan, A., Fruchard, L., Krin, E., Carvalho, A., Mazel, D., and Baharoglu, Z. (2022). Nonessential tRNA and rRNA modifications impact the bacterial response to sub-MIC antibiotic stress. Microlife 3, uqac019. 10.1093/femsml/uqac019.

62. Delhermite, J., Tafforeau, L., Sharma, S., Marchand, V., Wacheul, L., Lattuca, R., Desiderio, S., Motorin, Y., Bellefroid, E., and Lafontaine, D.L.J. (2022). Systematic mapping of rRNA 2’-O methylation during frog development and involvement of the methyltransferase Fibrillarin in eye and craniofacial development in Xenopus laevis. PLoS Genet. 18, e1010012. 10.1371/journal.pgen.1010012.

63. Fasnacht, M., Gallo, S., Sharma, P., Himmelstoß, M., Limbach, P.A., Willi, J., and Polacek, N. (2022). Dynamic 23S rRNA modification ho5C2501 benefits Escherichia coli under oxidative stress. Nucleic Acids Res. 50, 473–489. 10.1093/nar/gkab1224.

64. Barozzi, C., Zacchini, F., Corradini, A.G., Morara, M., Serra, M., De Sanctis, V., Bertorelli, R., Dassi, E., and Montanaro, L. (2023). Alterations of ribosomal RNA pseudouridylation in human breast cancer. NAR Cancer 5, zcad026. 10.1093/narcan/zcad026.

65. Zhang, M., Jiang, Z., Ma, Y., Liu, W., Zhuang, Y., Lu, B., Li, K., Peng, J., and Yi, C. (2023). Quantitative profiling of pseudouridylation landscape in the human transcriptome. Nat. Chem. Biol. 19, 1185–1195. 10.1038/s41589-023-01304-7.

66. Sklias, A., Cruciani, S., Marchand, V., Spagnuolo, M., Lavergne, G., Bourguignon, V., Brambilla, A., Dreos, R., Marygold, S.J., Novoa, E.M., et al. (2024). Comprehensive map of ribosomal 2’-O-methylation and C/D box snoRNAs in Drosophila melanogaster. Nucleic Acids Res. 52, 2848–2864. 10.1093/nar/gkae139.

67. Lin, R., Kos, A., Lopez, J.P., Dine, J., Fiori, L.M., Yang, J., Ben-Efraim, Y., Aouabed, Z., Ibrahim, P., Mitsuhashi, H., et al. (2023). SNORD90 induces glutamatergic signaling following treatment with monoaminergic antidepressants. Elife 12. 10.7554/eLife.85316.

68. Krogh, N., Jansson, M.D., Häfner, S.J., Tehler, D., Birkedal, U., Christensen-Dalsgaard, M., Lund, A.H., and Nielsen, H. (2016). Profiling of 2’-O-Me in human rRNA reveals a subset of fractionally modified positions and provides evidence for ribosome heterogeneity. Nucleic Acids Res. 44, 7884–7895. 10.1093/nar/gkw482.

69. Sharma, S., Marchand, V., Motorin, Y., and Lafontaine, D.L.J. (2017). Identification of sites of 2’-O-methylation vulnerability in human ribosomal RNAs by systematic mapping. Sci. Rep. 7, 11490. 10.1038/s41598-017-09734-9.

70. Jain, M., Olsen, H.E., Paten, B., and Akeson, M. (2016). The Oxford Nanopore MinION: delivery of nanopore sequencing to the genomics community. Genome Biol. 17, 239. 10.1186/s13059-016-1103-0.

71. Smith, M.A., Ersavas, T., Ferguson, J.M., Liu, H., Lucas, M.C., Begik, O., Bojarski, L., Barton, K., and Novoa, E.M. (2020). Molecular barcoding of native RNAs using nanopore sequencing and deep learning. Genome Res. 30, 1345–1353. 10.1101/gr.260836.120.

72. Djirackor, L., Halldorsson, S., Niehusmann, P., Leske, H., Capper, D., Kuschel, L.P., Pahnke, J., Due-Tønnessen, B.J., Langmoen, I.A., Sandberg, C.J., et al. (2021). Intraoperative DNA methylation classification of brain tumors impacts neurosurgical strategy. Neurooncol Adv 3, vdab149. 10.1093/noajnl/vdab149.

73. Vermeulen, C., Pagès-Gallego, M., Kester, L., Kranendonk, M.E.G., Wesseling, P., Verburg, N., de Witt Hamer, P., Kooi, E.J., Dankmeijer, L., van der Lugt, J., et al. (2023). Ultra-fast deep-learned CNS tumour classification during surgery. Nature 622, 842–849. 10.1038/s41586-023-06615-2.

74. Fafard-Couture, É., Bergeron, D., Couture, S., Abou-Elela, S., and Scott, M.S. (2021). Annotation of snoRNA abundance across human tissues reveals complex snoRNA-host gene relationships. Genome Biol. 22, 172. 10.1186/s13059-021-02391-2.

75. Genuth, N.R., and Barna, M. (2018). The Discovery of Ribosome Heterogeneity and Its Implications for Gene Regulation and Organismal Life. Mol. Cell 71, 364–374. 10.1016/j.molcel.2018.07.018.

76. Barna, M., Karbstein, K., Tollervey, D., Ruggero, D., Brar, G., Greer, E.L., and Dinman, J.D. (2022). The promises and pitfalls of specialized ribosomes. Mol. Cell 82, 2179–2184. 10.1016/j.molcel.2022.05.035.

77. Cozzuto, L., Delgado-Tejedor, A., Hermoso Pulido, T., Novoa, E.M., and Ponomarenko, J. (2023). Nanopore Direct RNA Sequencing Data Processing and Analysis Using MasterOfPores. Methods Mol. Biol. 2624, 185–205. 10.1007/978-1-0716-2962-8_13.

78. Lowe, T.M., and Eddy, S.R. (1999). A computational screen for methylation guide snoRNAs in yeast. Science 283, 1168–1171. 10.1126/science.283.5405.1168.

79. Schattner, P., Decatur, W.A., Davis, C.A., Ares, M., Jr, Fournier, M.J., and Lowe, T.M. (2004). Genome-wide searching for pseudouridylation guide snoRNAs: analysis of the Saccharomyces cerevisiae genome. Nucleic Acids Res. 32, 4281–4296. 10.1093/nar/gkh768.

80. Dobin, A., Davis, C.A., Schlesinger, F., Drenkow, J., Zaleski, C., Jha, S., Batut, P., Chaisson, M., and Gingeras, T.R. (2013). STAR: ultrafast universal RNA-seq aligner. Bioinformatics 29, 15–21. 10.1093/bioinformatics/bts635.

