## Supplementary figures for "Epitranscriptomic rRNA fingerprinting reveals tissue-of-origin and tumor-specific signatures"

**Figure S1. Replicability plots of SE values along the 18S rRNA, across tissues and developmental stages.** Scatterplots depicting the summed base-calling errors values of two biological replicates across mouse tissues and developmental stages.

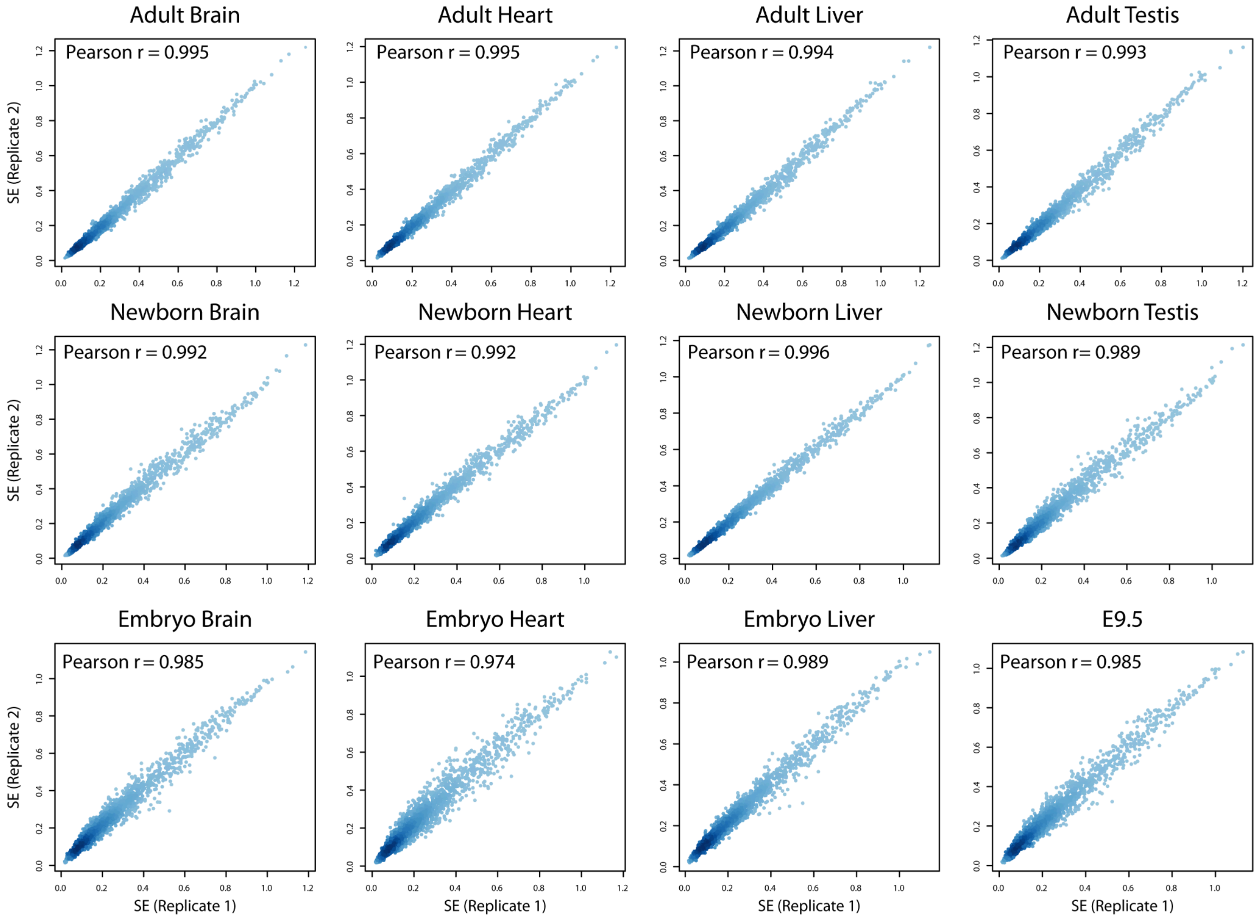

**Figure S2. Replicability plots of SE values along the 28S rRNA, across tissues and developmental stages.** Scatterplots depicting the summed base-calling errors values of two biological replicates across mouse tissues and developmental stages.

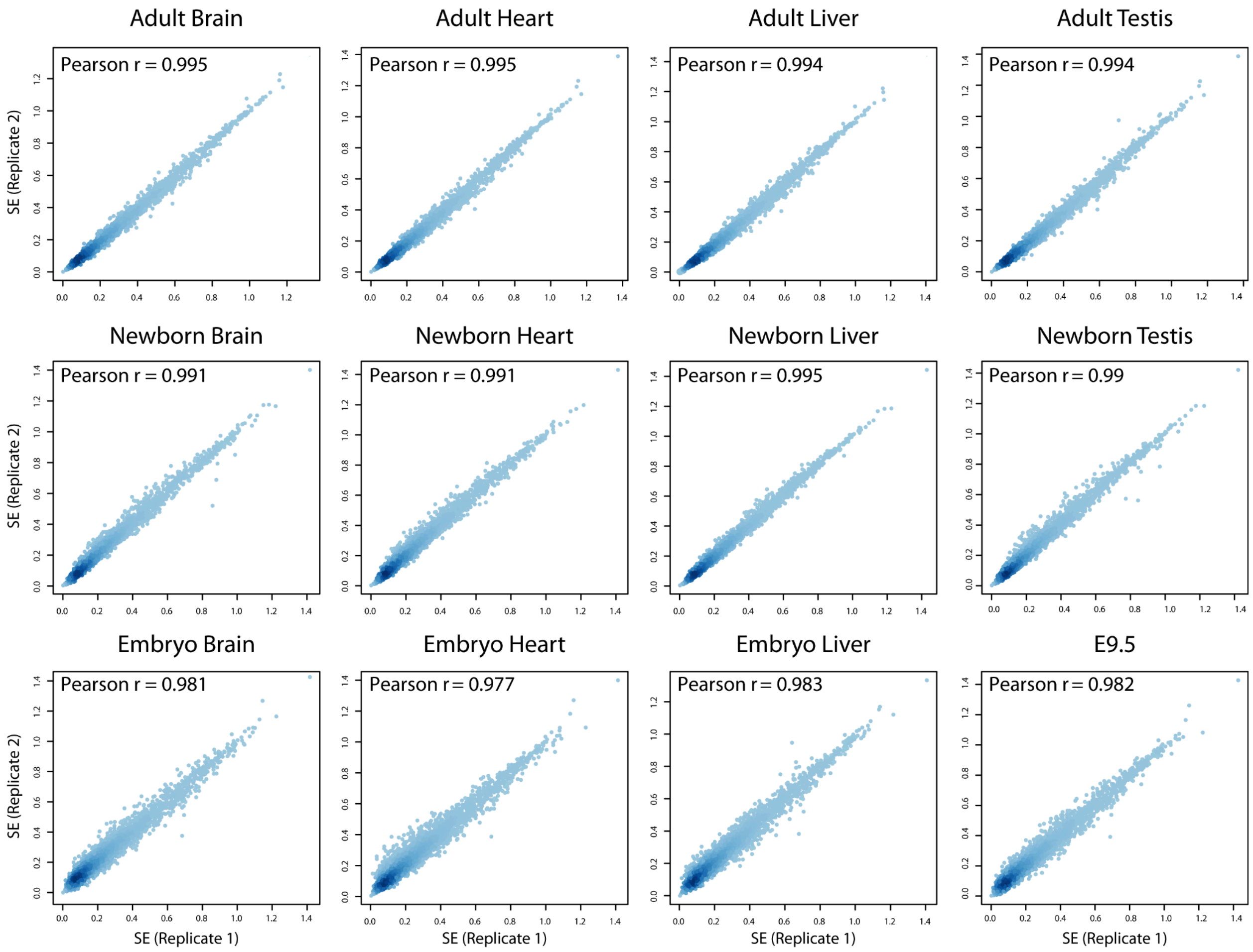

**Figure S3. Identification of sample-specific RNA modifications in 18S rRNA (Replicate 1).** Scatterplots depicting the summed base-calling errors values of a given sample, relative to the median of summed basecalling error values from all tissues and developmental stages.

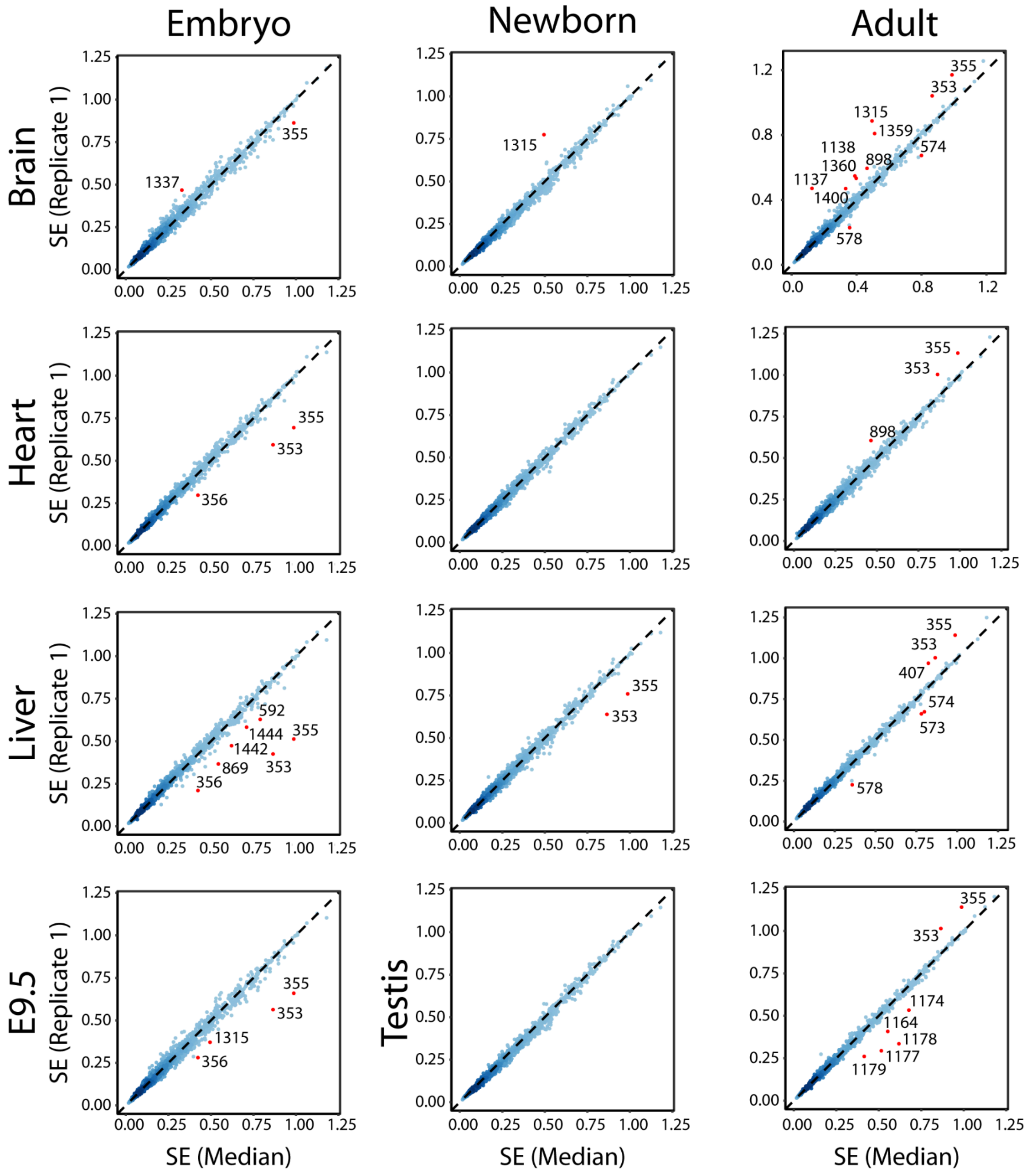

**Figure S4. Identification of sample-specific RNA modifications in 18S rRNA (Replicate 2).** Scatterplots depicting the summed base-calling errors values of a given sample, relative to the median of summed basecalling error values from all tissues and developmental stages.

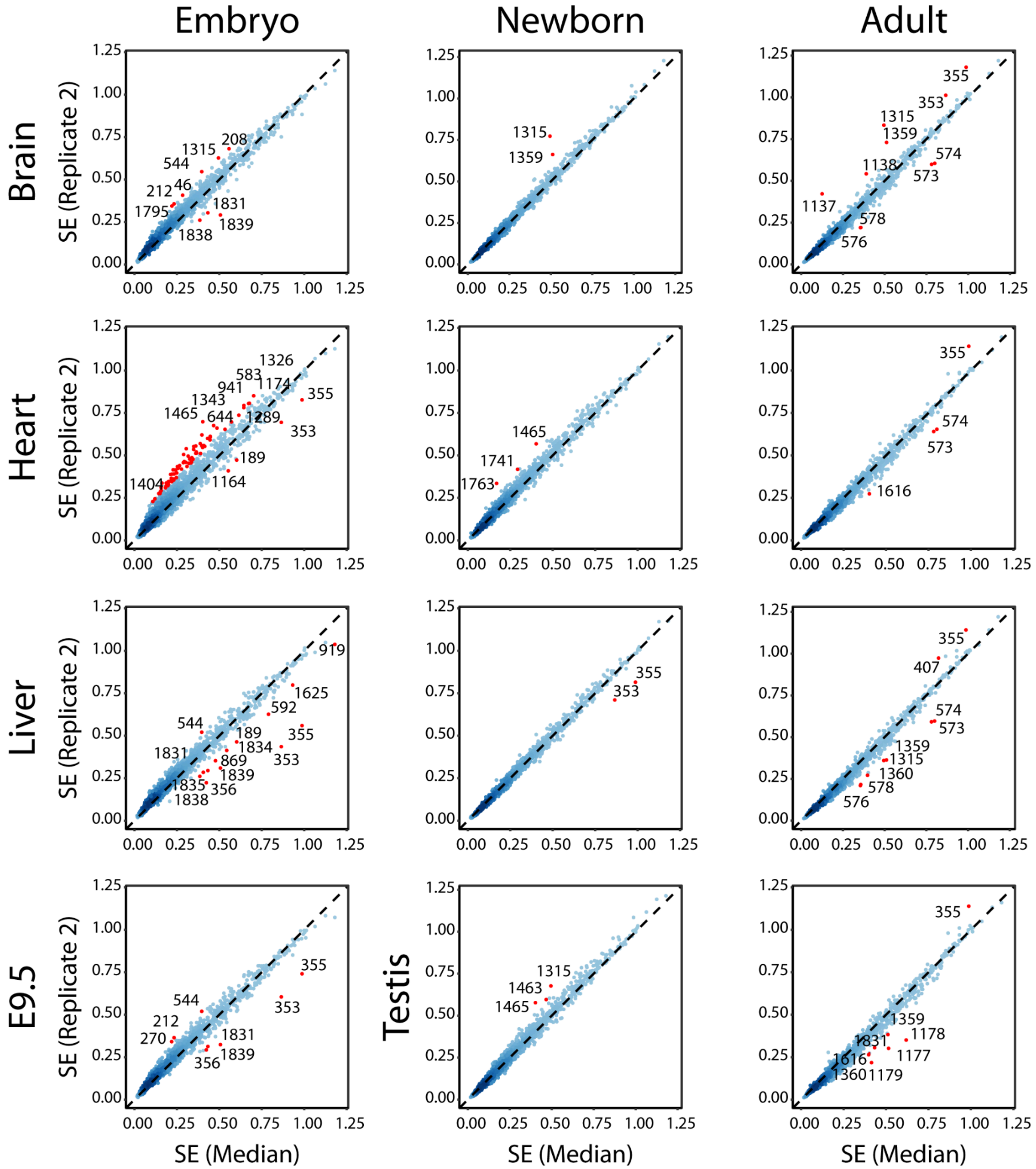

**Figure S5. Identification of sample-specific RNA modifications in 28S rRNA (Replicate 1).** Scatterplots depicting the summed base-calling errors values of a given sample, relative to the median of summed basecalling error values from all tissues and developmental stages.

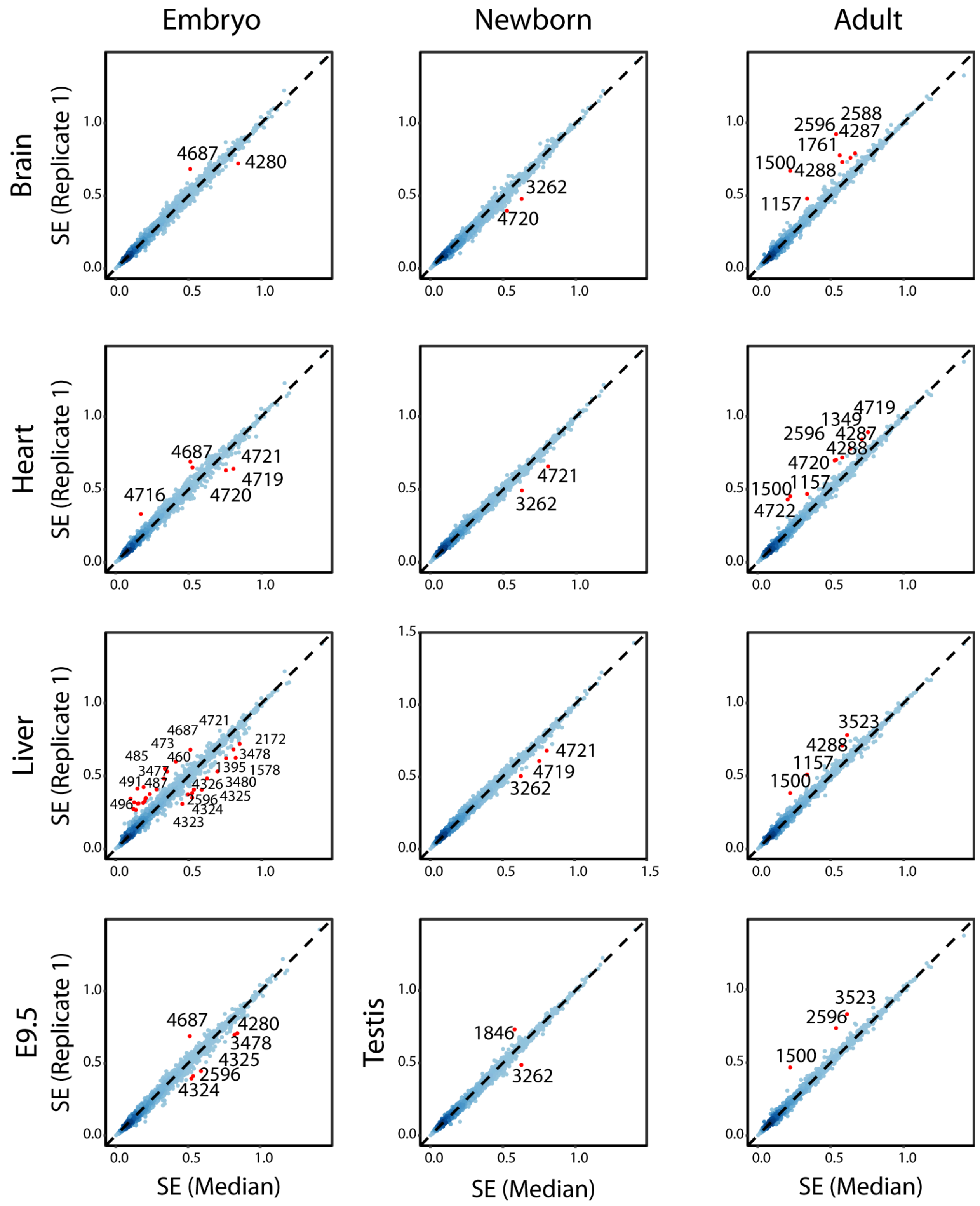

**Figure S6. Identification of sample-specific RNA modifications in 28S rRNA (Replicate 2).** Scatterplots depicting the summed base-calling errors values of a given sample, relative to the median of summed basecalling error values from all tissues and developmental stages.

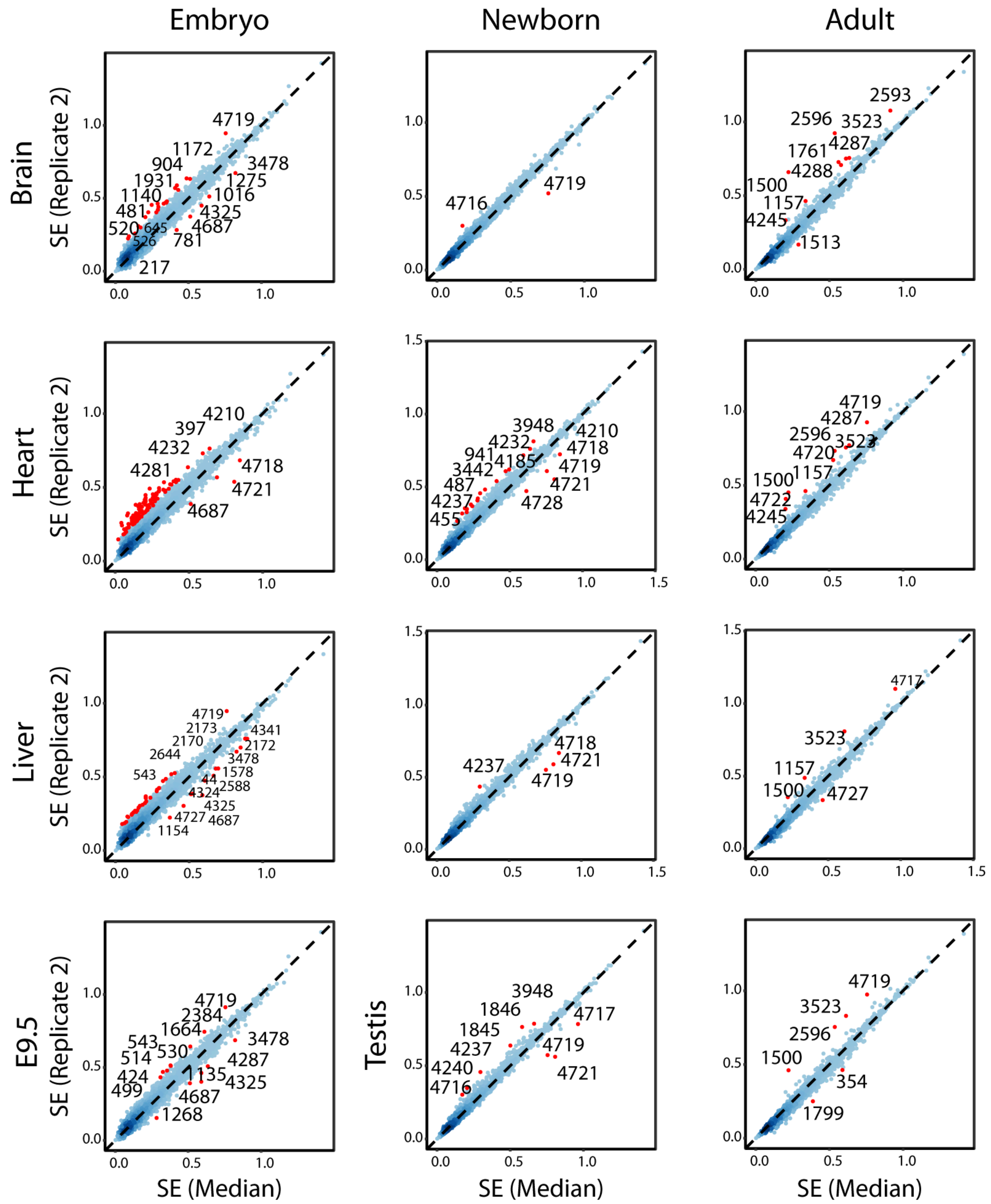

**Figure S7. Summed errors of all annotated rRNA modification sites in mouse tissues.** Heatmap depicting the summed base-calling errors of previously annotated rRNA modification sites across mouse tissues and developmental stages.

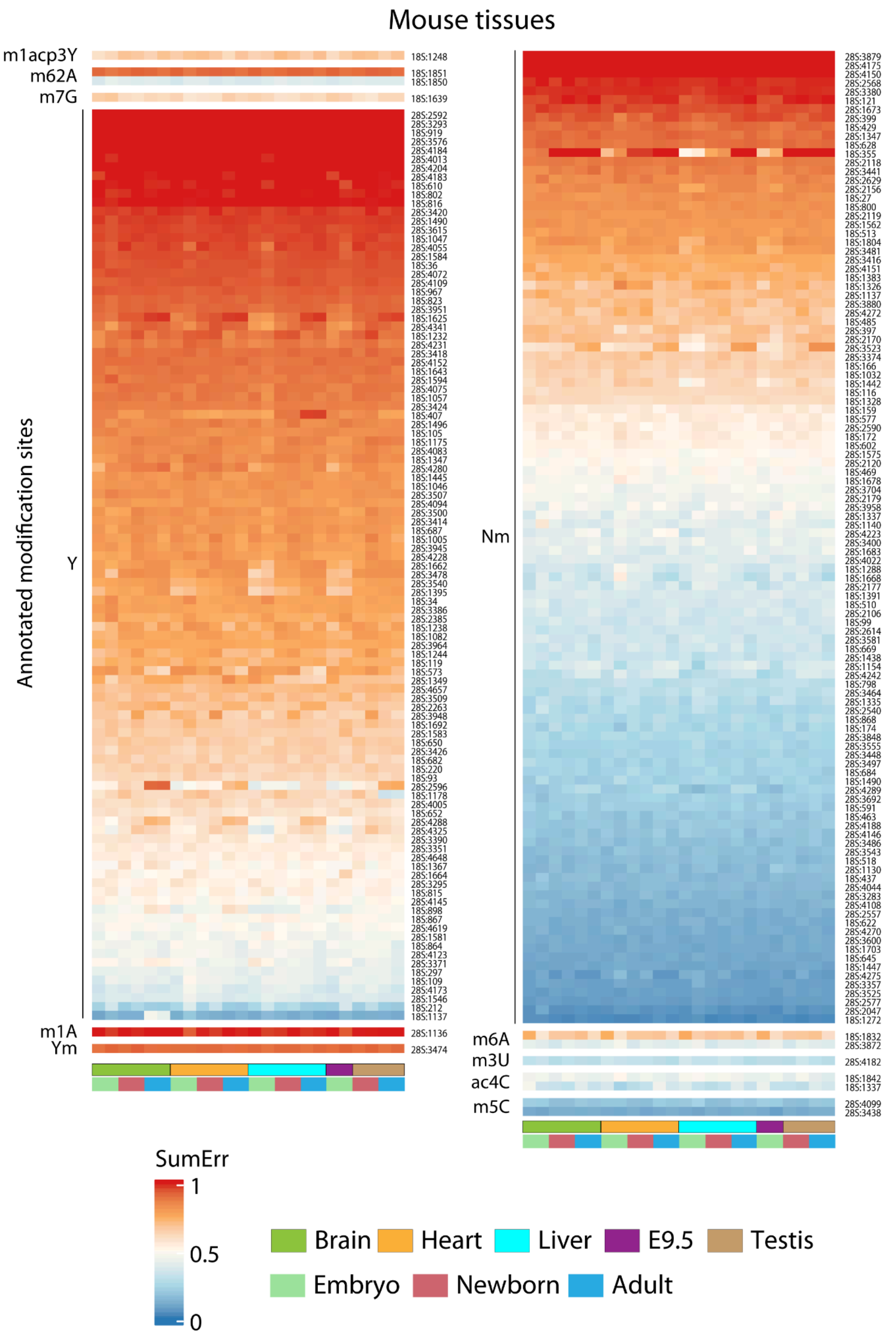

**Figure S8. NanoCMC validation of putative pseudouridine sites in 18S.** Peaks correspond to previously annotated or newly discovered pseudouridylation sites. Dashed lines indicate the CMC score threshold used for determining the pseudouridylation sites.

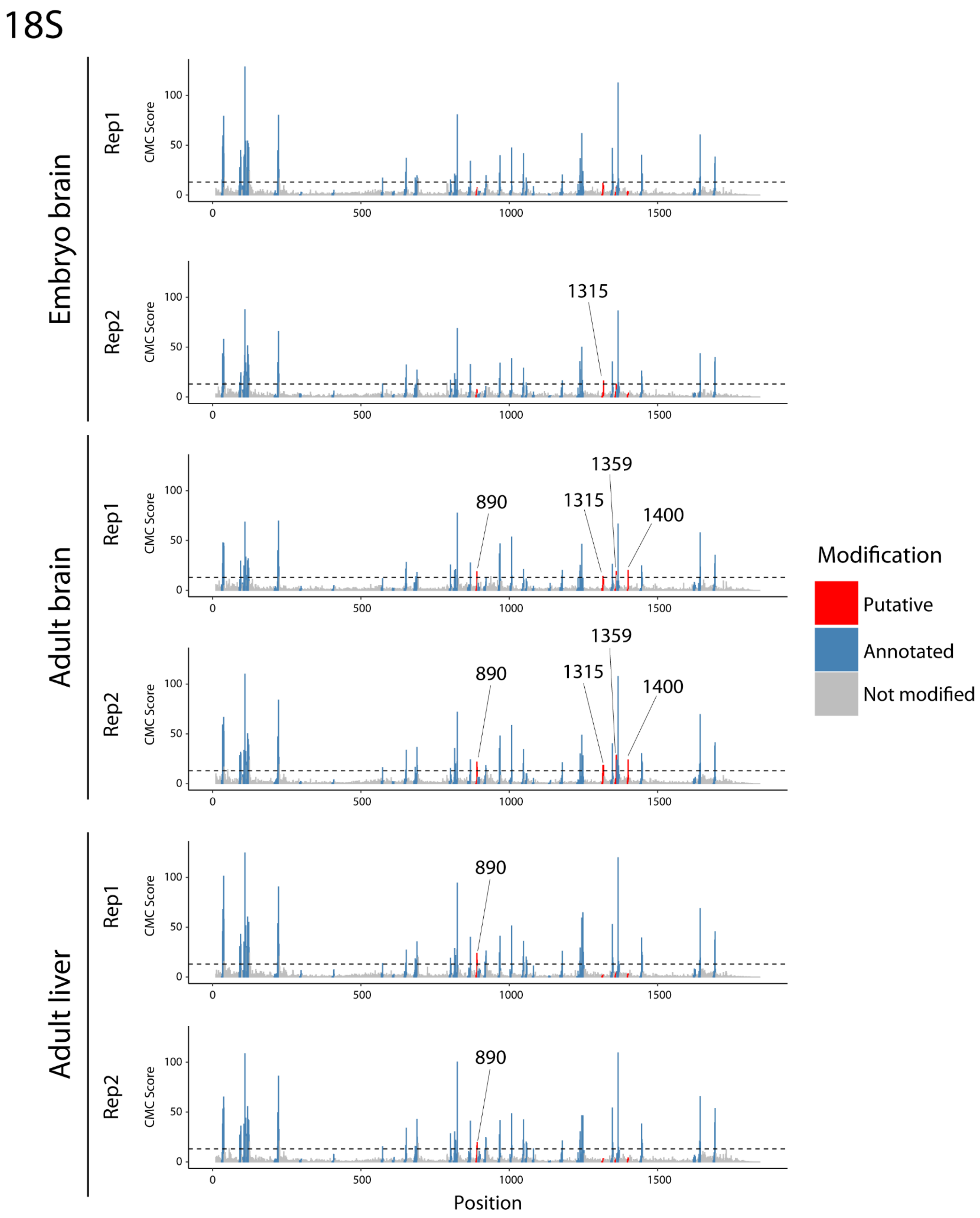

**Figure S9. NanoCMC validation of putative pseudouridine sites in 28S.** Peaks correspond to previously annotated or newly discovered pseudouridylation sites. Dashed lines indicate the CMC score threshold used for determining the pseudouridylation sites.

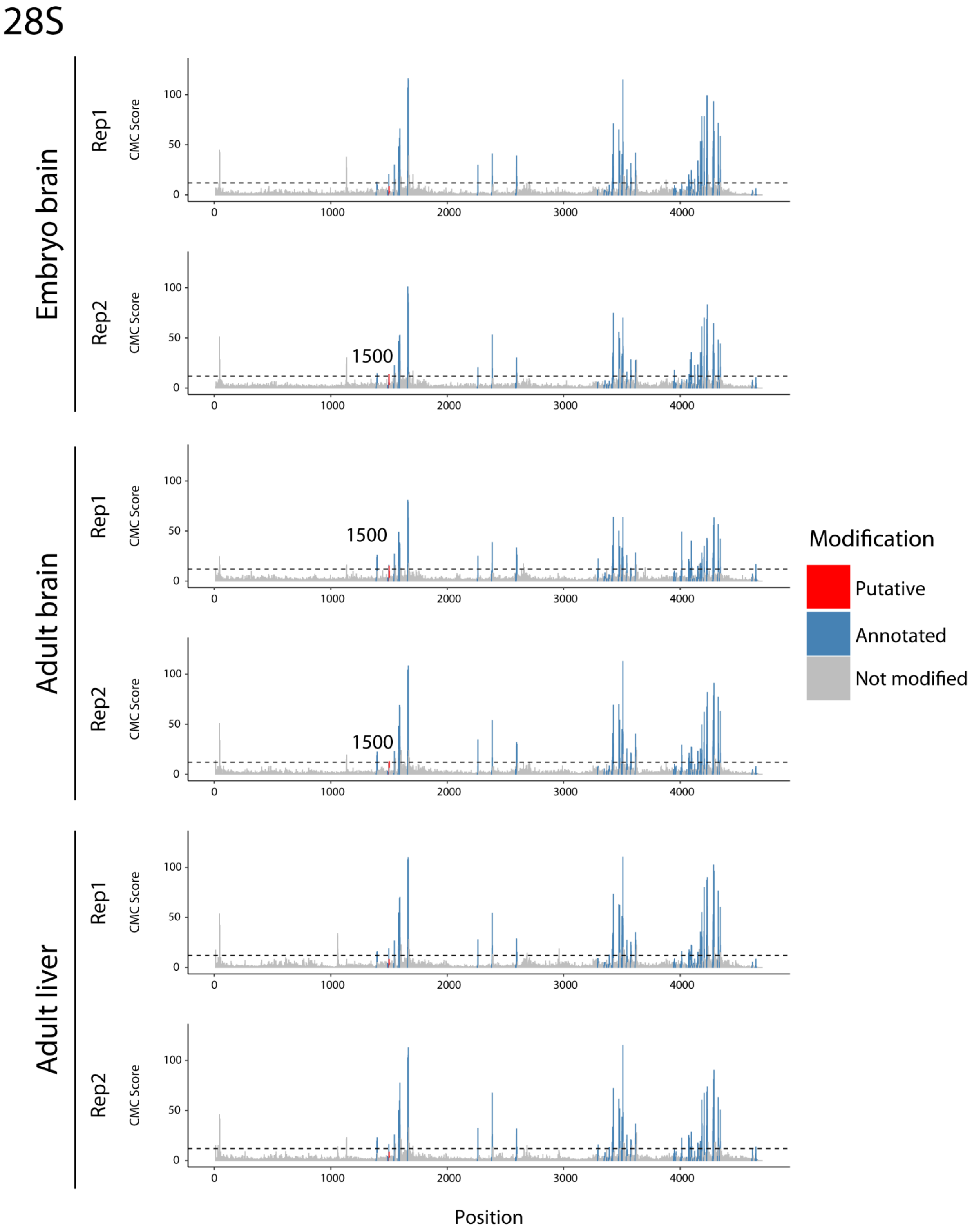

**Figure S10. snoRNA predictions for three unannotated dynamic rRNA modification sites. (A)** *Snora35b* and 18S:1315 duplex. **(B)** *Snora-Taf1* and 18S:1359 duplex. **(C)** *Snora-Wwc1* and 28S:1500 duplex.

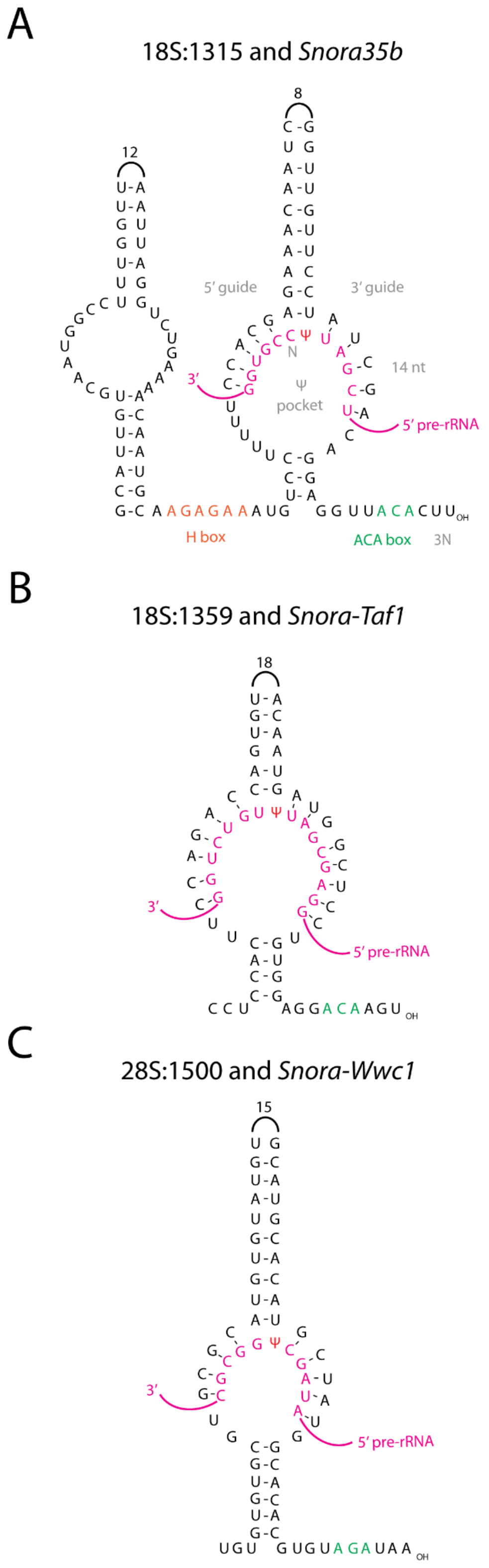

**Figure S11. rRNA modification levels and snoRNA modification levels for some of the annotated dynamic modification sites. (A)** rRNA modification levels (top, expressed in SE) and snoRNA expression levels (bottom, CPM) for four annotated modification sites with known snoRNAs. **(B)** rRNA modification levels (expressed in SE) for four annotated modification sites with unknown snoRNAs. The snoRNA expression data was obtained from Isakova et al. [1].

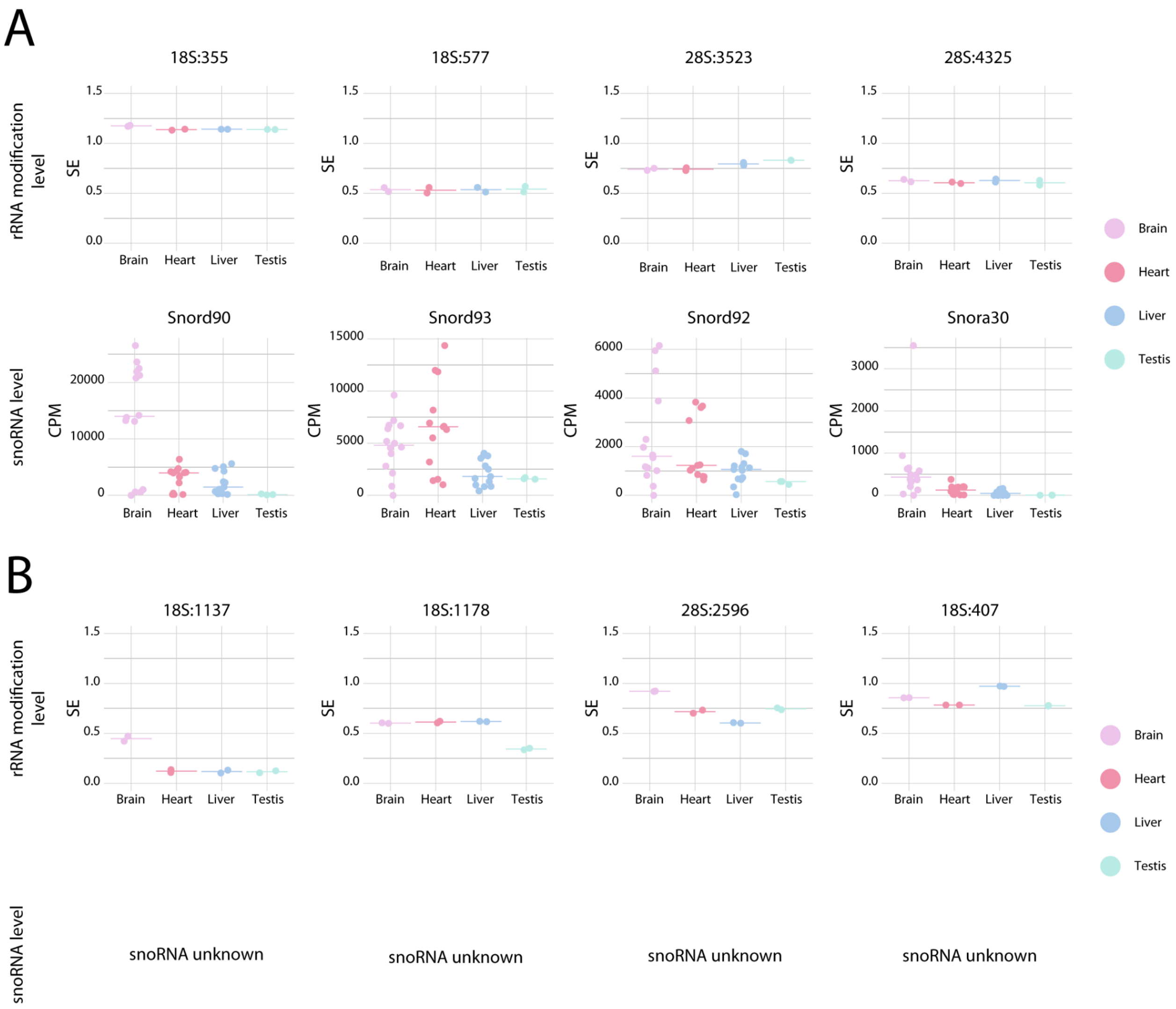

**Figure S12. Identification of sample-specific RNA modifications in neurons and mESc in 18S rRNA.** Scatterplots depicting the summed base-calling errors values of a given sample, relative to the median of summed basecalling error values from all tissues and developmental stages.

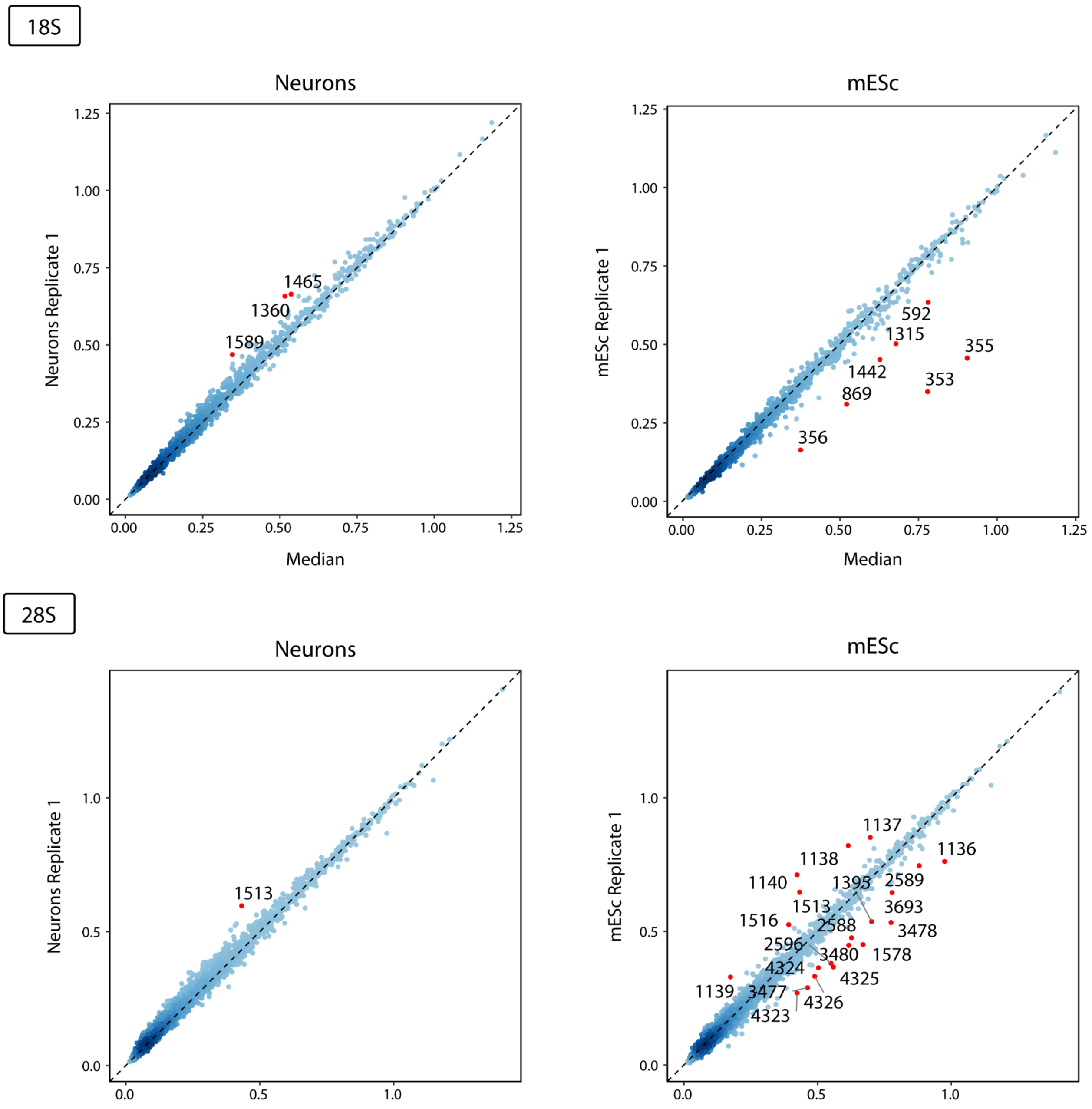

**Figure S13. Summed errors of all annotated rRNA modification sites in mouse brain and cells.** Heatmap depicting the summed base-calling errors of previously annotated rRNA modification sites in the embryonic, newborn and adult mouse brain and mES, NPC and neuronal cells.

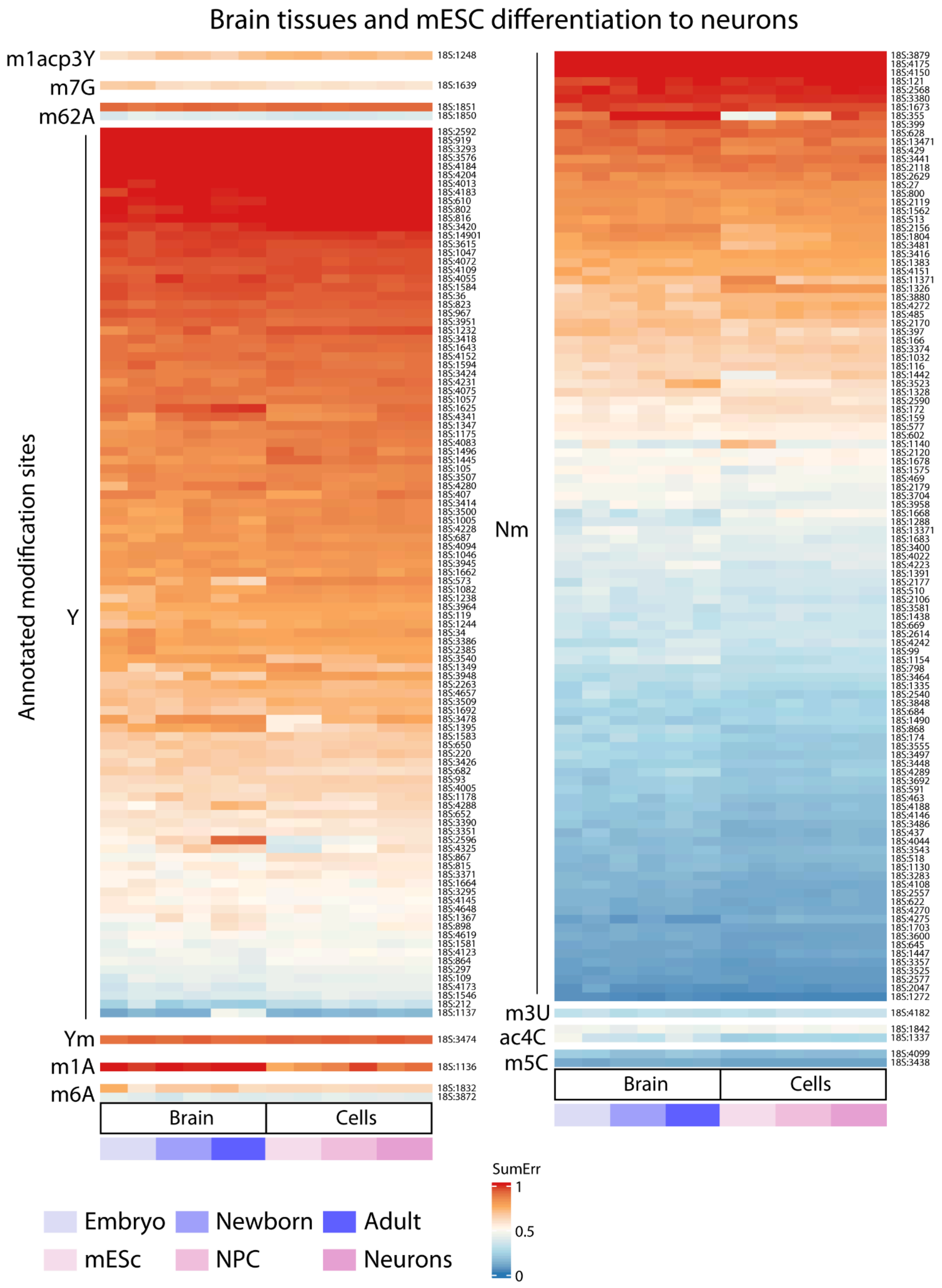

**Figure S14. Workflow used for the training, testing and validating the RF tissue classifier. (A)** Schematic representation of the approach used for training a tissue classifier. (**B**) Performance of the RF classifier on the independent validation set, which was sequenced in a third independent flowcell (not used for training nor testing).

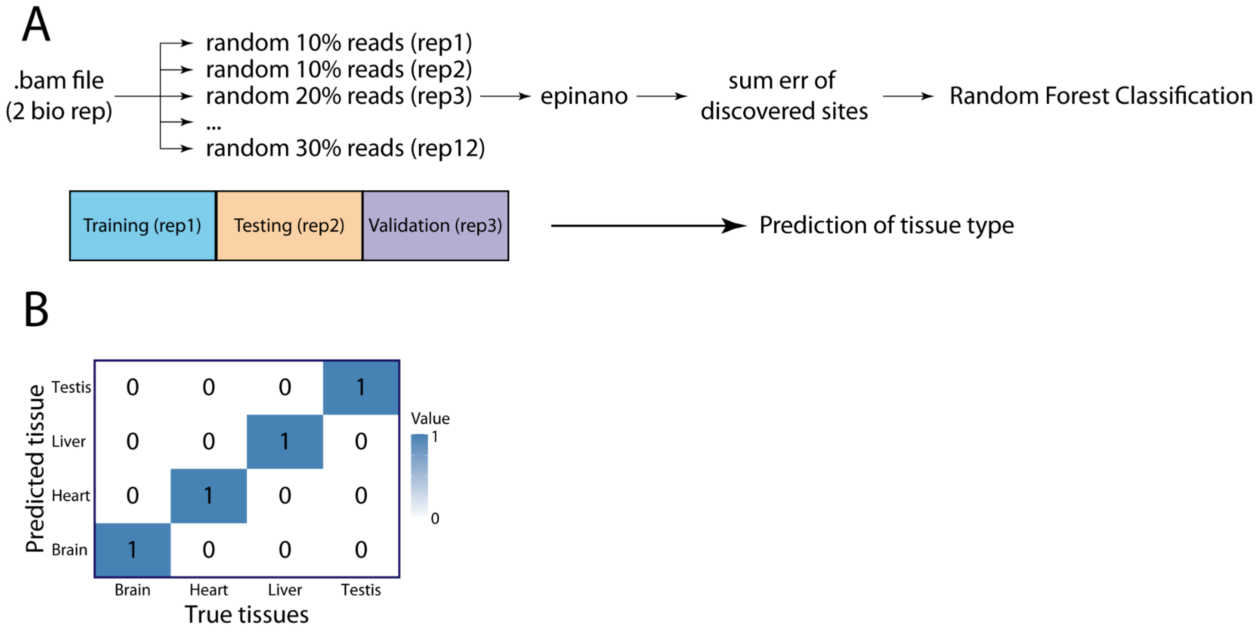

**Figure S15. Workflow used for the training and testing of the RF cell type classifier and its performance. (A)** Schematic representation of the approach used for training a cell type classifier. (**B**) Performance of the RF classifier on the testing data set (training set was sequenced in an independent flowcell and the reads were subdivided into 12 pseudoreplicates).

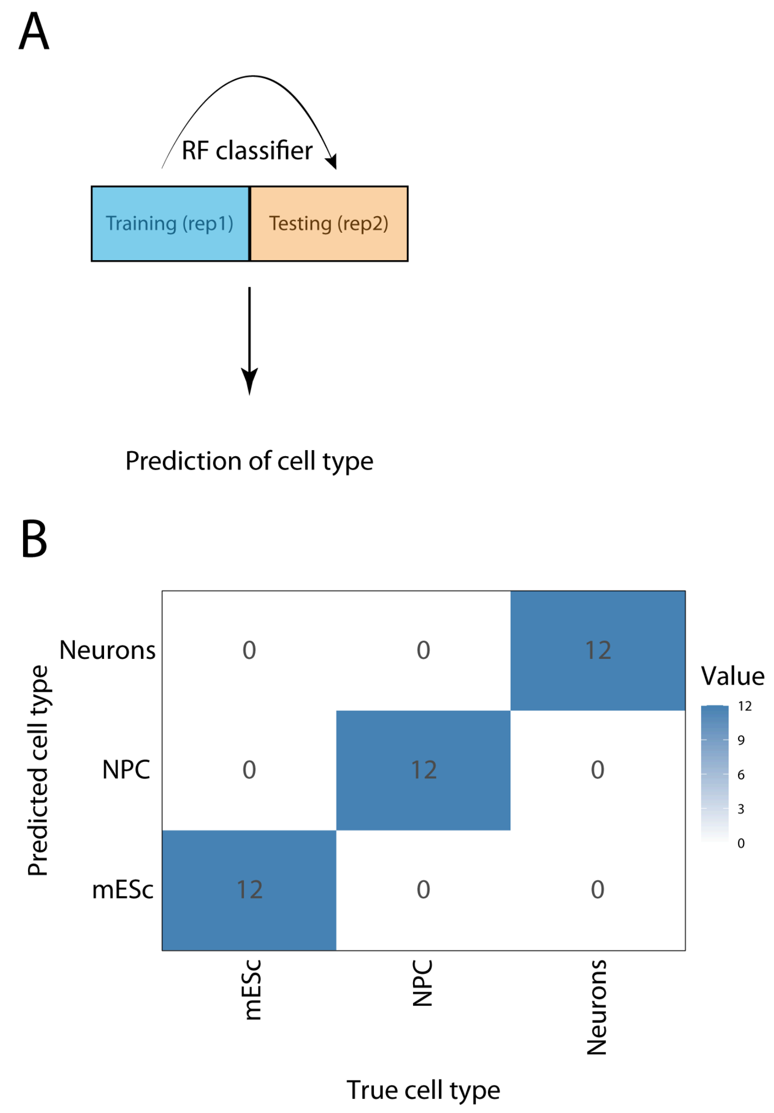

**Figure S16. Workflow used for the training and testing of the RF tissue stage classifier and its performance. (A)** Schematic representation of the approach used for training a tissue classifier. (**B**) Performance of the RF classifier on the testing data set (training set was sequenced in an independent flowcell and the reads were subdivided into 12 pseudoreplicates).

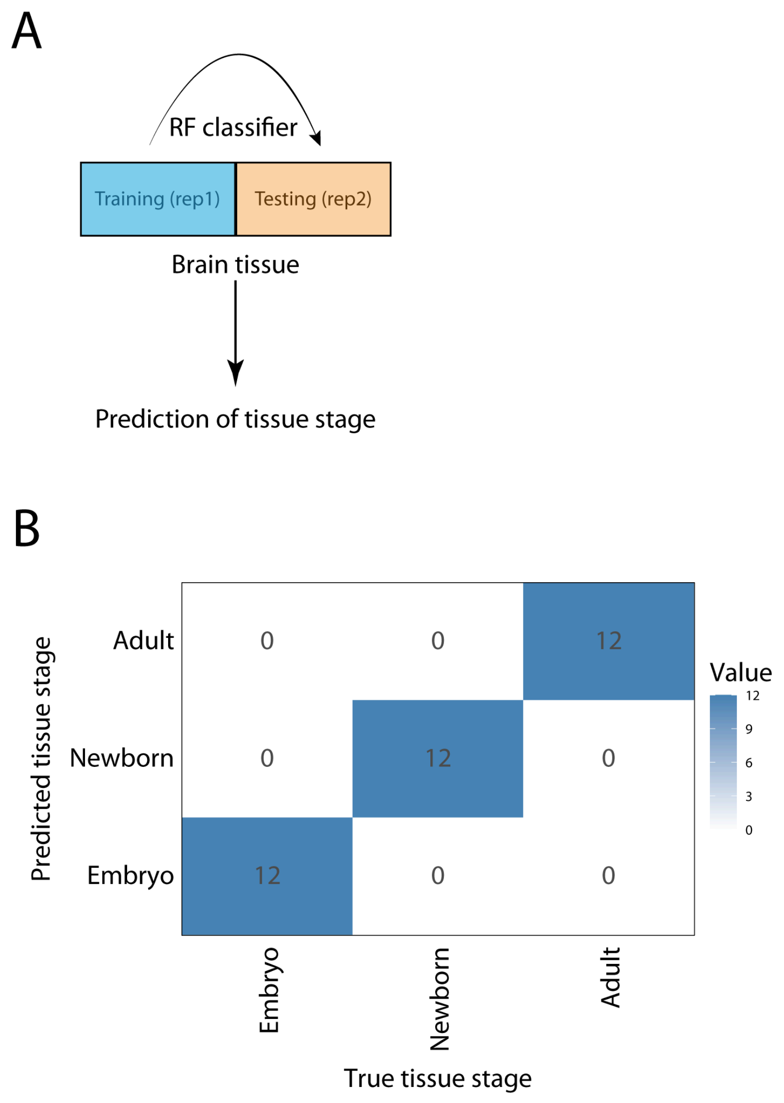

**Figure S17. PCA of mouse tissues performed on SE values of dynamic rRNA modification sites.** Each tissue is shown in a different shape, whereas each stage is shown in the form of a different color. The variance explained by each principal component is also shown in the axis.

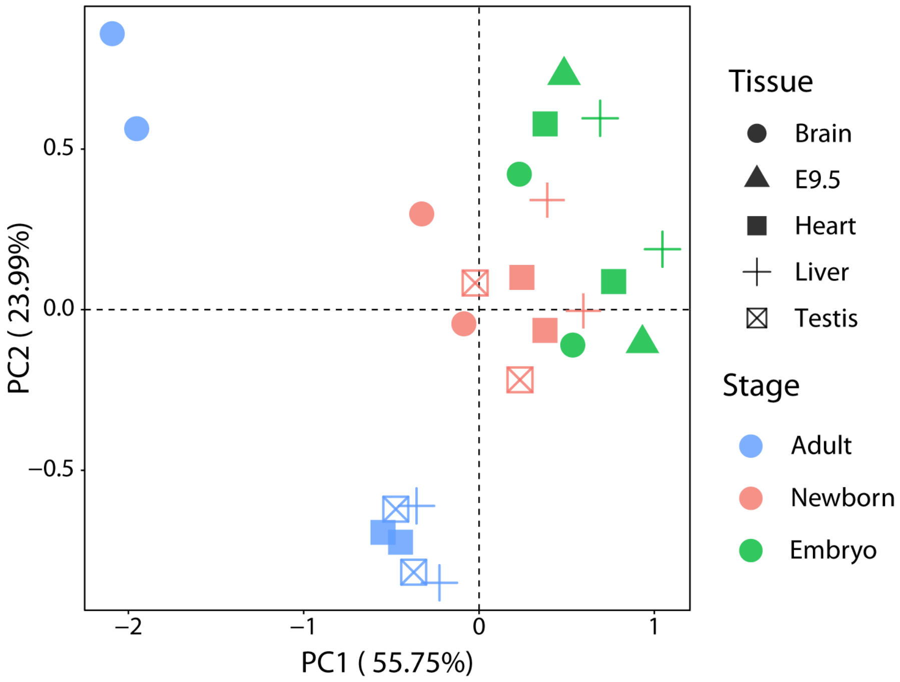

**Figure S18. Difference in summed errors along the 18s and 28s rRNA transcripts, comparing the WT and Snord90KO cells.** The 18S:Um355 position is highlighted in red (and depicted by an arrow), and corresponds to the largest difference in summed errors between WT and Snord90 KO cells. We should note that this position is also present in mESC, but the difference between WT and Snord90KO is smaller (lower ΔSE), suggesting that this position is modified at lower stoichiometry in mESC, compared to neurons.

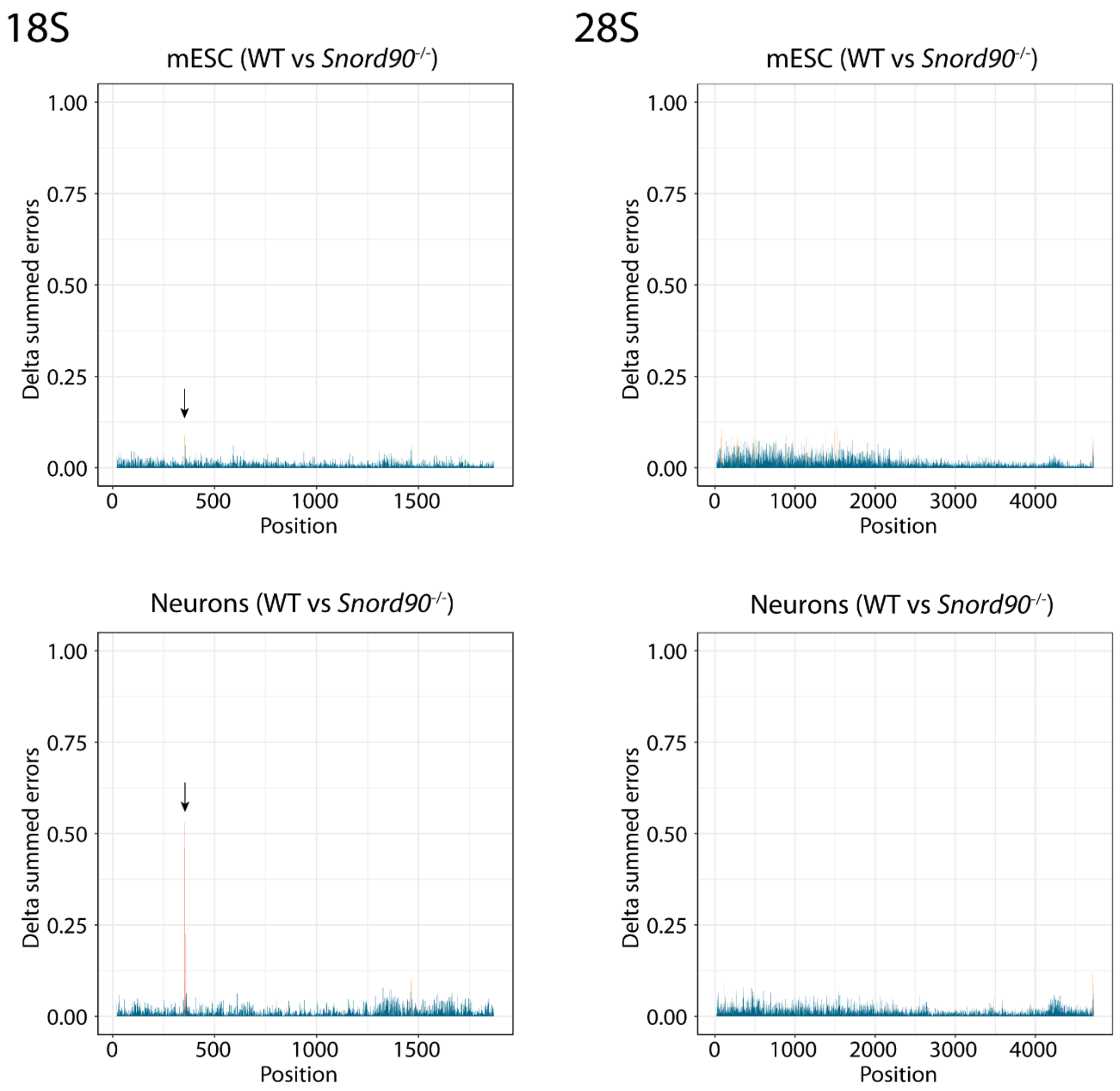

**Figure S19. Training and testing a RF classifier for lung cancer prediction.** (**A**) Scheme of the workflow used for classifier training and testing. (**B**) Performance of the classifier on the testing dataset.

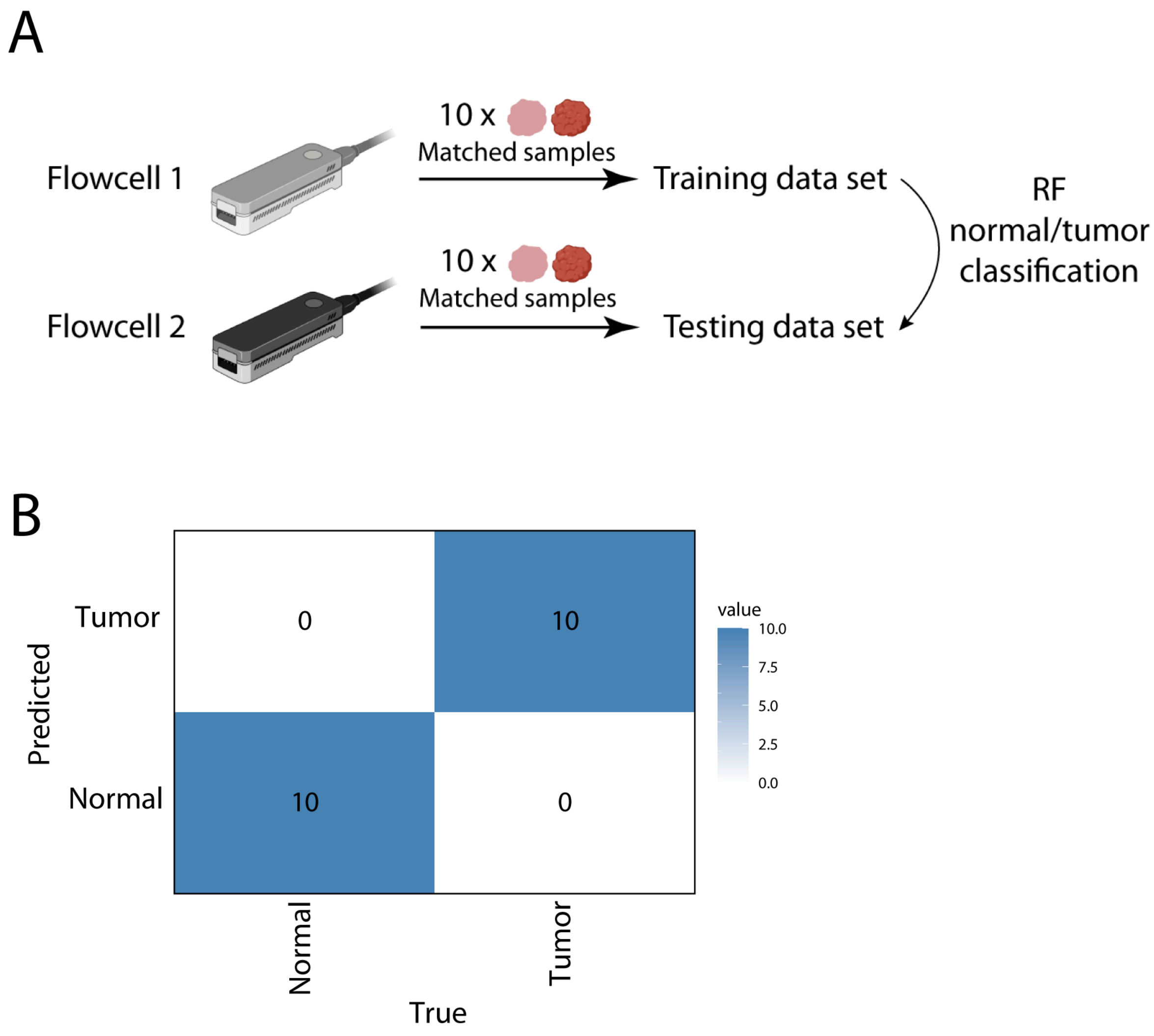

**Figure S20. PCA done using the cancer metadata. (A)** PCA of cancer samples based on the tumor stage (**I** and **II**), calculated on the top 20 dynamic sites between the two stages. **(B)** PCA of the cancer samples, based on whether the patients developed metastasis, calculated on the top 20 dynamic sites between metastatic and non-metastatic samples (see also **Table S11** for metadata for each patient and sample).

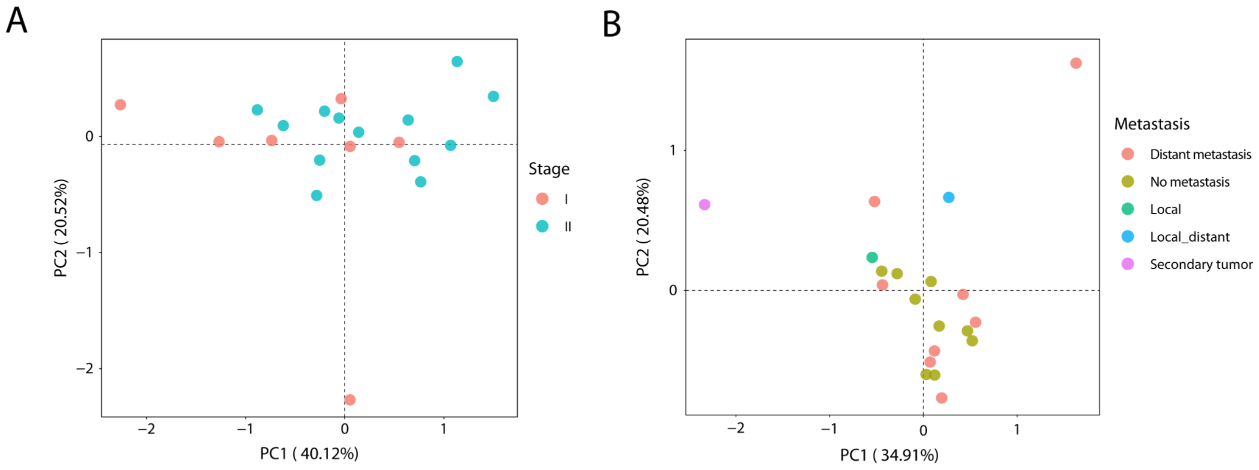

**Figure S21. Normal-tumor separation and classification based on different subsets of dynamic rRNA sites.** (**A**) Principal component analysis showing the separation of normal and tumor matched lung cancer samples, using as input the SE values of the top 20 dynamic rRNA sites, when considering annotated Nm modifications (left panel), annotated Ψ modified sites (middle panel) and both annotated and unannotated sites (the 'agnostic' approach, right panel). (**B**) ROC curve performance of the Random Forest classifier using as input the top 20 dynamic annotated Nm sites (left), annotated Ψ modified sites (middle) and both annotated and unannotated sites (right), when using a varying number of reads as input. The area under the curve (AUC) is shown for each number of input reads, for each panel.

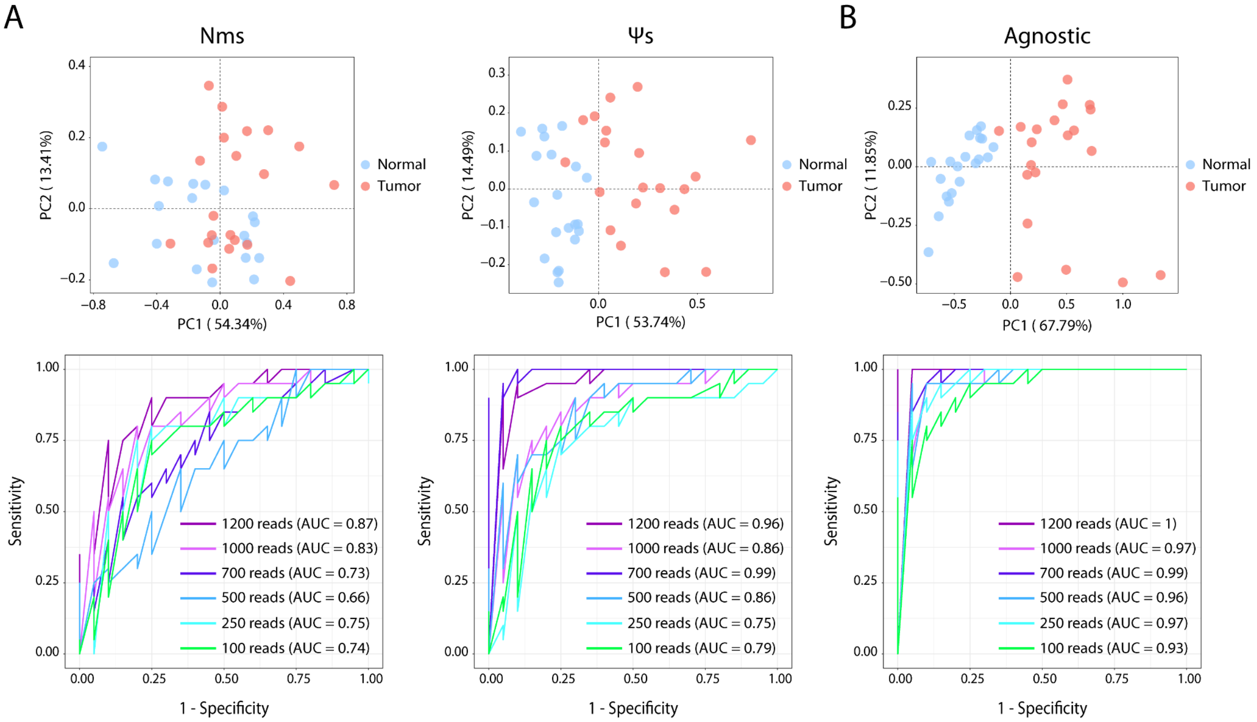

**Figure S22. Unannotated putative modification sites in mouse 18S and 28S rRNA.** IGV tracks showing the previously unannotated dynamic putative modification sites across cells and mouse tissues discovered using direct RNA nanopore sequencing.

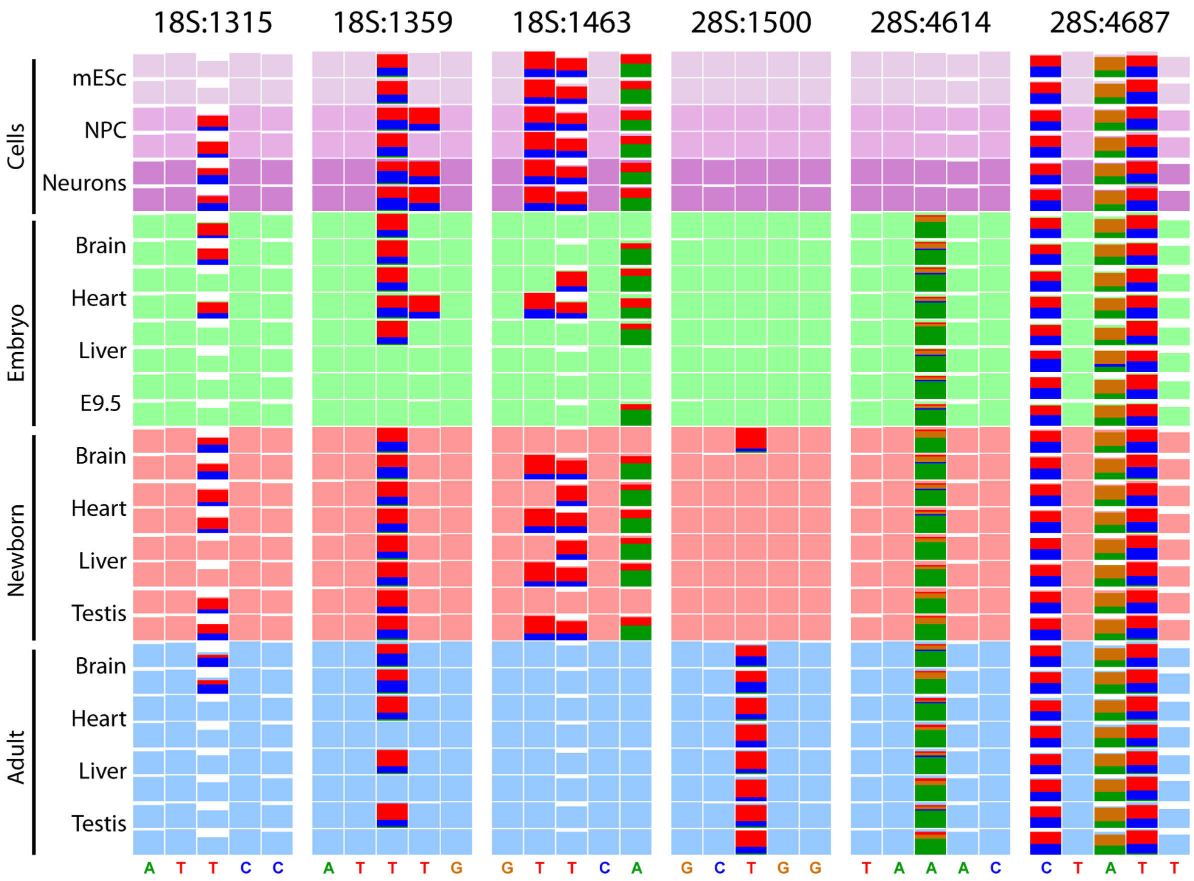

**Figure S23. Tapestation total RNA profiles of matched normal-tumor lung samples.** Each sample and ID corresponds to a different patient. The upper lanes correspond to total RNA profiles from ‘normal’ tissue samples, and the bottom total RNA profiles correspond to ‘tumor’ samples. The RIN values are shown below each lane, for each sample.

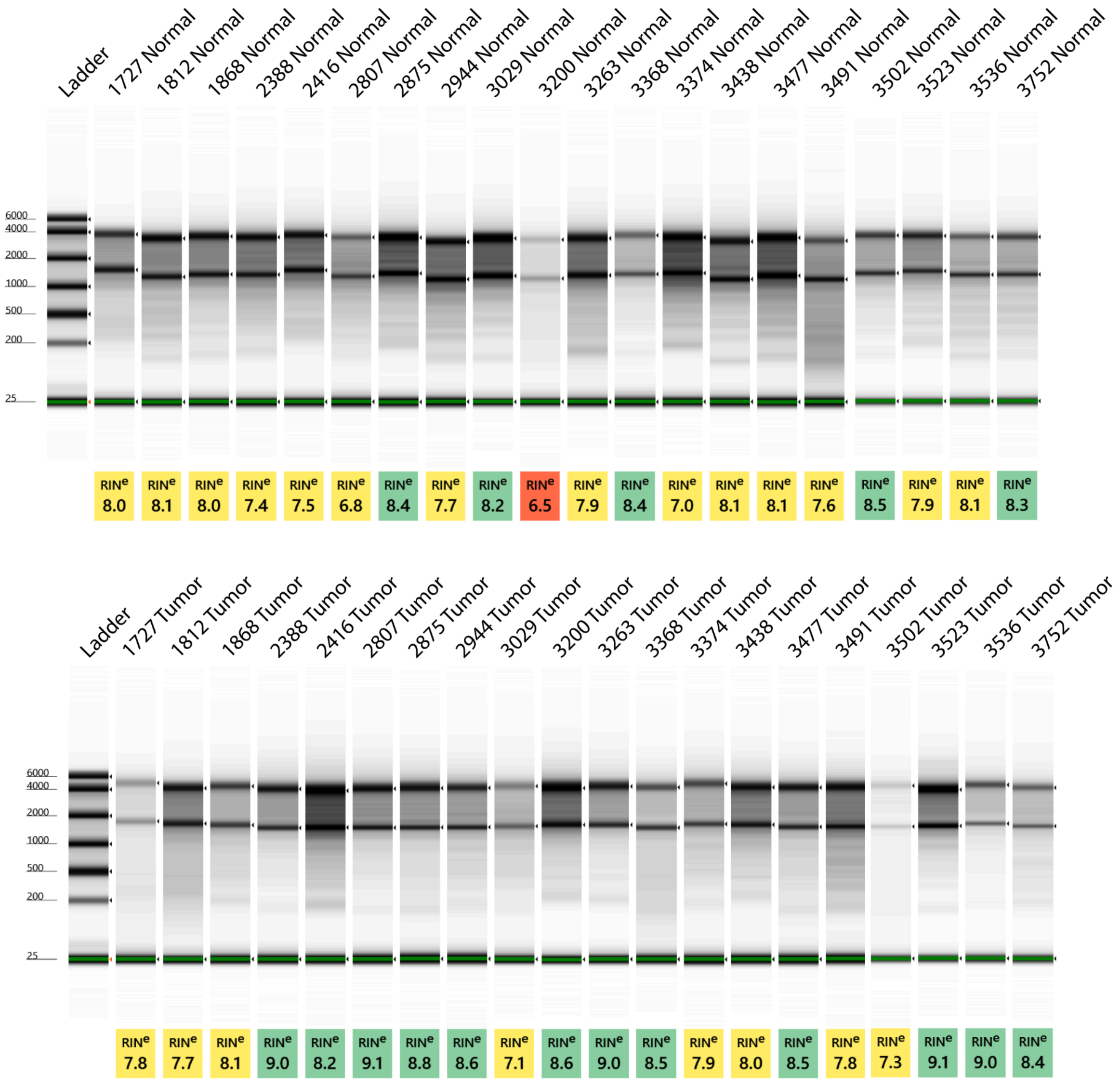

### SUPPLEMENTARY REFERENCES

1. Isakova, A., Fehlmann, T., Keller, A., and Quake, S.R. (2020). A mouse tissue atlas of small noncoding RNA. Proc. Natl. Acad. Sci. U. S. A. *117*, 25634–25645.
